# Automated evaluation of single-cell reference atlas mappings enables the identification of disease-associated cell states

**DOI:** 10.1101/2025.05.23.655749

**Authors:** Lisa Sikkema, Mariam Lkandushi, Daniele Scarcella, Amir Ali Moinfar, Jan Engelmann, Fabian J. Theis

## Abstract

The rise of single-cell atlases has opened up the use of reference atlases as a comprehensive healthy control for the study of disease. A key step for using atlases is the mapping of “query” disease or perturbed datasets onto a healthy reference atlas, which allows for comparison of the query to healthy controls and thereby the identification of perturbation-specific transcriptional cell states. However, the mapping success, i.e. the extent to which these mappings correctly reflect biological similarities and differences between query and reference, varies with e.g. experimental confounders in the data (dissociation protocol, single-cell assay) or the mapping method used, and cannot be predicted in advance. Moreover, an unsuccessful mapping can result in falsely reporting technical artifacts as disease or perturbation-specific cellular changes and in overlooking true altered states. Here, we present MapQC, a method that quantifies query-to-reference mapping success by leveraging the information present in the reference embedding. Specifically, MapQC tests the presence of remaining batch effect in the embedding after mapping by comparing inter-sample distances in the healthy reference itself to distance of query control samples to the reference. Similarly, it tests whether disease or perturbation-specific variation has been retained during mapping by testing whether query perturbed samples are more distant to the reference than reference samples are from each other. We apply MapQC to lung and endometrial query datasets, including data of idiopathic pulmonary fibrosis and Asherman Syndrome, mapped to large-scale healthy references. We show that MapQC correctly distinguishes successful mappings from unsuccessful ones where data visualization techniques such as UMAP can be deceiving, and that mapQC outperforms integration metrics often re-purposed for mapping qualitycontrol. Moreover, only mappings that MapQC identified as successful result in the correct identification of cell state changes specific to disease. Taken together, MapQC provides a critical quality-control metric for a more reliable, robust, and accurate use of reference atlases to study disease.

## 1 Introduction

Large-scale cellular reference atlases are being constructed for a variety of tissues and organs[1], and are used increasingly widely for the annotation and interpretation of new data [2–4]. Reference atlases bring together data from many different studies and thereby offer a uniquely broad view of the phenotypic variation existing in a population, including among healthy individuals. Whereas single datasets often only include a limited set of healthy controls, complicating the distinction between normal variation and disease-induced changes, healthy tissue atlases offer a more comprehensive collection of healthy controls. Thus, comparing newly generated data of disease or other perturbations (e.g. genetic knock-outs, drug treatments, etc.) to an existing healthy reference atlas enables a more accurate distinction between “normal” and perturbation-specific variation [5, 6], an essential goal of biomedical research.

Several methods have been developed to allow for a fast and easy comparison of newly generated data (i.e. a “query”) to an existing atlas (i.e. a “reference”) [7–10]. These methods enable the mapping of query data onto an existing low-dimensional representation of a reference, for example using transfer learning[8, 11]. The resulting joint representation can be used for the interpretation of the query data, such as the identification of cell types from diseased tissue in the query data that are different from their healthy counterparts [5, 6].

For the correct interpretation of query data mapped onto a reference atlas, it is essential that the variation captured in the mapped data reflects the true biological variation. If, instead, variation caused by technical covariates (e.g. sequencing platform or tissue dissociation protocol) is preserved in the mapping, disease-specific effects are likely to be confounded with technical artifacts in the data, making it impossible to confidently infer new disease-specific knowledge from the query-to-reference comparison without prior knowledge. Conversely, if a mapping removes too much variation from the query, disease-specific changes can become undetectable. Metrics to capture and quantify these possible mapping failures are therefore crucial.

A number of metrics are currently used to characterize the quality of query-to-reference mappings. One group of metrics measures the preservation of variation in the query before and after mapping, for example by quantifying preservation of query cell neighborhoods or clusters[9, 10, 12]. These metrics require that the topology of the query’s embedding remains unchanged after mapping, which defeats the purpose of the mapping - namely, that by using the reference during embedding, the query’s topology changes, revealing subtle sources of variation in the query such as rare cell types. Moreover, the metrics are insensitive to the extent to which the query and the reference mix, an essential aspect of mapping quality. A second group of metrics consists of general batch integration metrics, that are now re-purposed to assess the quality of a mapping by treating the reference as one batch and the query as another. For example, some metrics quantify the mixing of cells from distinct donors or batches in the latent space after integration or mapping [9, 13]. Whereas such metrics are useful as relative metrics, i.e. to compare different mapping approaches applied to the same data in a benchmark setting, they fail to define clear thresholds to distinguish a failed from a successful mapping and are therefore difficult to interpret. Most importantly, none of these metrics take into account the fact that there is not just one healthy state, but that variation exists even within the healthy population. Acknowledging and leveraging this fact opens the path to a new and better way to evaluate query-to-reference mappings. Here we present MapQC, a metric to assess the success of the mapping of a query dataset onto an existing healthy reference atlas, independent of the mapping method used. Leveraging the comprehensive coverage of healthy phenotypes in a reference, MapQC learns the inter-sample distances expected between healthy samples at the cell-state level. It then validates successful removal of batch effects by performing a control test on the query control samples, checking whether query control sample distances to the reference samples are as expected based on the reference. Finally, MapQC ensures that biological variation has been retained during mapping by performing a case test, in which it checks whether case samples from the query (e.g. from diseased tissue or perturbed cells) display a larger distance to the reference. We apply MapQC to two query datasets mapped to two distinct references, and show that MapQC identifies mappings in which batch effects were not successfully removed. We furthermore demonstrate that such mapping failures, if left undetected, could lead to false identification of putative disease effects. In contrast, mappings identified as successful by MapQC, e.g. through improved mapping parameter settings, enable the correct identification of disease-specific cell states. Overall, given the upcoming wide use of reference atlases for the analysis and interpretation of new data, MapQC is an essential tool for guiding correct and reliable atlas use.

## 2 Results

### 2.1 MapQC uses local intersample distances to evaluate mappings

To distinguish a successful from an unsuccessful query-to-reference mapping, MapQC builds on two basic assumptions:

#### Assumption 1

Healthy reference atlases capture most of the cellular variation present within the healthy population.

#### Assumption 2

Diseased tissue is transcriptionally different from healthy tissue for one or more cell types.

From these assumptions, two hypotheses can be derived that should hold true in the case of a successful query-to-reference mapping. These are the two hypotheses that MapQC tests for a given mapping.

#### Hypothesis 1

Healthy “control” samples in the query are equally similar to healthy samples in the reference, as healthy reference samples are similar to each other.

#### Hypothesis 2

“Case” samples (of e.g. diseased individuals) are, at least for some cell types, less similar to healthy reference samples, than healthy reference samples are similar to each other.

Based on hypothesis 1 and 2, we can establish data-driven thresholds to distinguish between successful and failed mappings, with query control samples similar to the reference and query case samples dissimilar to the reference in the case of a successful mapping (**fig. 1a**). In this framework, dissimilar samples are defined as those whose distance from the reference falls outside of the normal range as observed in the healthy reference. MapQC thus uniquely leverages a key aspect of healthy reference atlases: given their wide coverage of the healthy population, they enable learning what normal, healthy inter-sample variation looks like.

**Figure 1.**
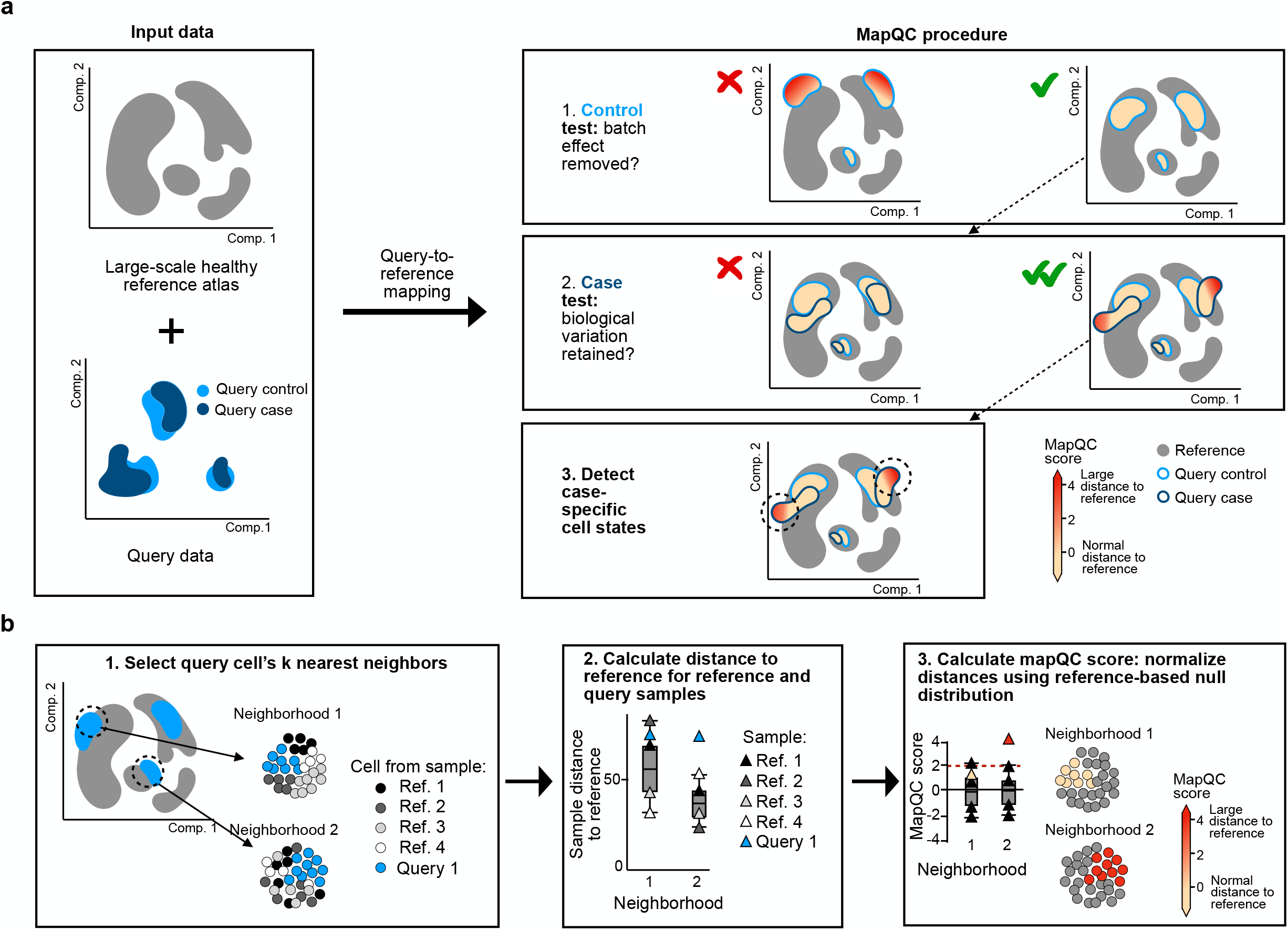
MapQC evaluates query-to-reference mapping quality, enabling correct identification of case (e.g. disease)-specific cell states. **a**, MapQC workflow. Left: mapQC can be used when mapping a dataset (“query data”) that includes control samples, and optionally “case” samples (e.g. diseased or perturbed) onto a large-scale reference atlas of control samples. Right: The MapQC procedure includes two tests: first, it tests whether batch effects were sufficiently removed during the mapping by testing if the query mixes with the reference. If distance to the reference is large for control cells, the mapping has failed. Second, if case (e.g. disease)-specific states are expected to be present, it tests whether biological variation has been retained by checking whether some of the cells from the case samples do not mix with the reference, indicating a casespecific cell state. If both tests are passed, downstream analysis can be performed and case-specific cell states can be identified and analyzed. **b**, Calculation of mapQC scores. First (1) neighborhoods of size k around randomly sampled query cells are calculated. Then (2) within each neighborhood, the mean distance to the reference samples is calculated for both query and reference samples. Finally (3), the distances are normalized per neighborhood, such that reference sample distances to the reference have mean 0 and standard deviation 1 in each neighborhood. Query cell distances to the reference are considered large if they are more than 2 standard deviations above the reference mean (dashed red line), in which case the mapQC score is higher than 2. These cells can be considered as not mixing with the reference.

Concretely, to establish the “null distribution” of normal inter-sample variation as observed in the reference, distances between reference samples themselves are first calculated (including only control samples from the reference). These distances are calculated in the integrated embedding space using the Energy distance metric[14, 15] (**Methods**). Notably, normal inter-sample distances can differ from cell type to cell type and even from state to state. To account for this cell-type specific level of inter-sample variation, baselines are calculated within large neighborhoods of cells instead of on the entire atlas (**fig. 1b**). Indeed, inter-sample distances in the reference vary markedly between neighborhoods, even within the same cell type, as do their variances (**Extended Data fig. 1**), supporting the use of local inter-sample distance null-distributions over global. Taken together, a sample’s distance to the reference *d_h_*(*s_i_, ref*) in a given neighborhood *h* is calculated as its mean distance to all reference samples in that neighborhood:

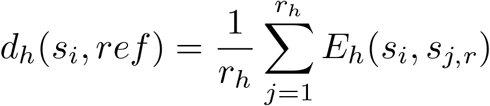

where *r_h_* is the number of samples from the reference in neighborhood *h* that have passed filtering (see **Methods**), *s_j,r_* is a sample from the reference, *s_i_* ≠ *s_j_*, and *E_h_*(*s_i_, s_j_*) is the energy distance between *s_i_* and *s_j_*. The distances to the reference for the set of reference samples in a given neighborhood together form that neighborhood’s null distribution.

Using the neighborhood-specific null-distributions, sample distances to the reference can now be normalized across neighborhoods, such that a sample’s normalized distance - in a given neighborhood-is a measure of its distance to the reference as compared to inter-sample distances in the reference itself (**fig. 1b**). Normalizing distances accordingly results in a standardized and interpretable measure of a sample’s distance to the reference across neighborhoods, and even across mappings. The normalization is performed by Z-scoring distances to the reference within neighborhoods, using the neighborhood’s null distributions mean and standard deviation for the Z-scoring, such that a sample *s_i_*’s normalized distance to the reference in a given neighborhood *h* is calculated as follows:

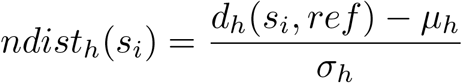

where *s_i_* is sample i, and *µ_h_* and *σ_h_* are the mean and standard deviation of the reference samples’ own distance to the reference, as based on the neighborhood’s null distribution. A cell *c_i_*’s mapQC score is then calculated as its sample *s*’s mean normalized distance to the reference across all the neighborhoods that the cell was part of (*N_c_*):

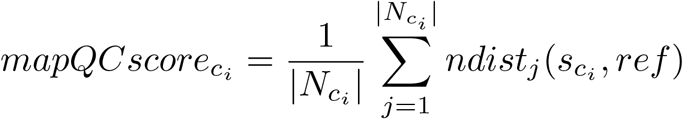

This results in a cell-level resolution score indicating for each cell whether it is close to or distant from the reference. A mapQC score of 0 for a cell means that in the cell’s neighborhood(s), its sample has a distance to the reference equal to the mean intersample distances in the reference itself, indicating high similarity of the sample to the reference. A mapQC score above 2 means a sample’s distance to the reference is more than two standard deviations above the reference’s mean inter-sample distance (which is expected for approximately 2% of samples from the null distribution, assuming it is Gaussian), indicating dissimilarity to the reference. We thus define a mapQC score*>*2 for a query cell as likely indicating either remaining batch effect or the presence of biological states not observed in the reference (fig. 1a).

MapQC’s cell-level resolution (each query cell has its own mapQC score) enables mapping analysis at any resolution of choice. In practice, as outliers are often expected, we recommend analyzing mapQC scores initially at the aggregate level, e.g. looking at the score distribution per cluster or cell type. This can aid in identifying groups of cells consistently mapping distant from the reference.

The mapQC scores can then used to determine if a query-to-reference mapping was successful, or if it has failed in one of two possible ways: by under-integrating or by over-integrating the query data into the reference (**fig. 1a**). A mapping was successful if query control samples are close to the reference (mapQC score*<*2) across the mapping, indicating no remaining batch effect, while query case samples have subset of cells distant to the reference (mapQC score*>*2), indicating preservation of case-specific cell states. If this is the case, the mapping can safely be used for downstream analysis. In contrast, the mapping has failed due to under-integration if distances between control samples in the query dataset and the healthy reference are larger than expected (mapQC score*>*2) based on the baselines, such that query-specific batcheffect is likely still present in the embedding. Conversely, the mapping has failed due to over-integration if case-related variation, such as a disease-related shift in gene expression, has been removed from the embedding, such that case samples consistently look equally similar to healthy reference samples as healthy reference samples are similar to each other.

### 2.2 MapQC identifies remaining batch effects after mapping

To investigate whether mapQC evaluates mappings correctly, we applied it to the mapping of an IPF dataset[16] including 5 samples from patients with IPF, and 6 control samples, onto the Human Lung Cell Atlas[6] (**fig. 2a, Extended Data fig. 2a, Methods**). First, to evaluate the presence of remaining batch effect in the query after mapping, we looked at mapQC scores for the control samples of the query (**fig. 2a-c, Extended Data fig. 2b**, query controls). Many cells were distant from the reference (mapQC-score*>*2), suggesting batch effects were still strongly present in the data (**fig. 2b, c**, query controls). Notably, this could not be visually inferred from the UMAP embedding in all cases (e.g. blood vessel endothelium, alveolar epithelium **fig. 2a, b, Extended Data fig. 2a**).

**Figure 2.**
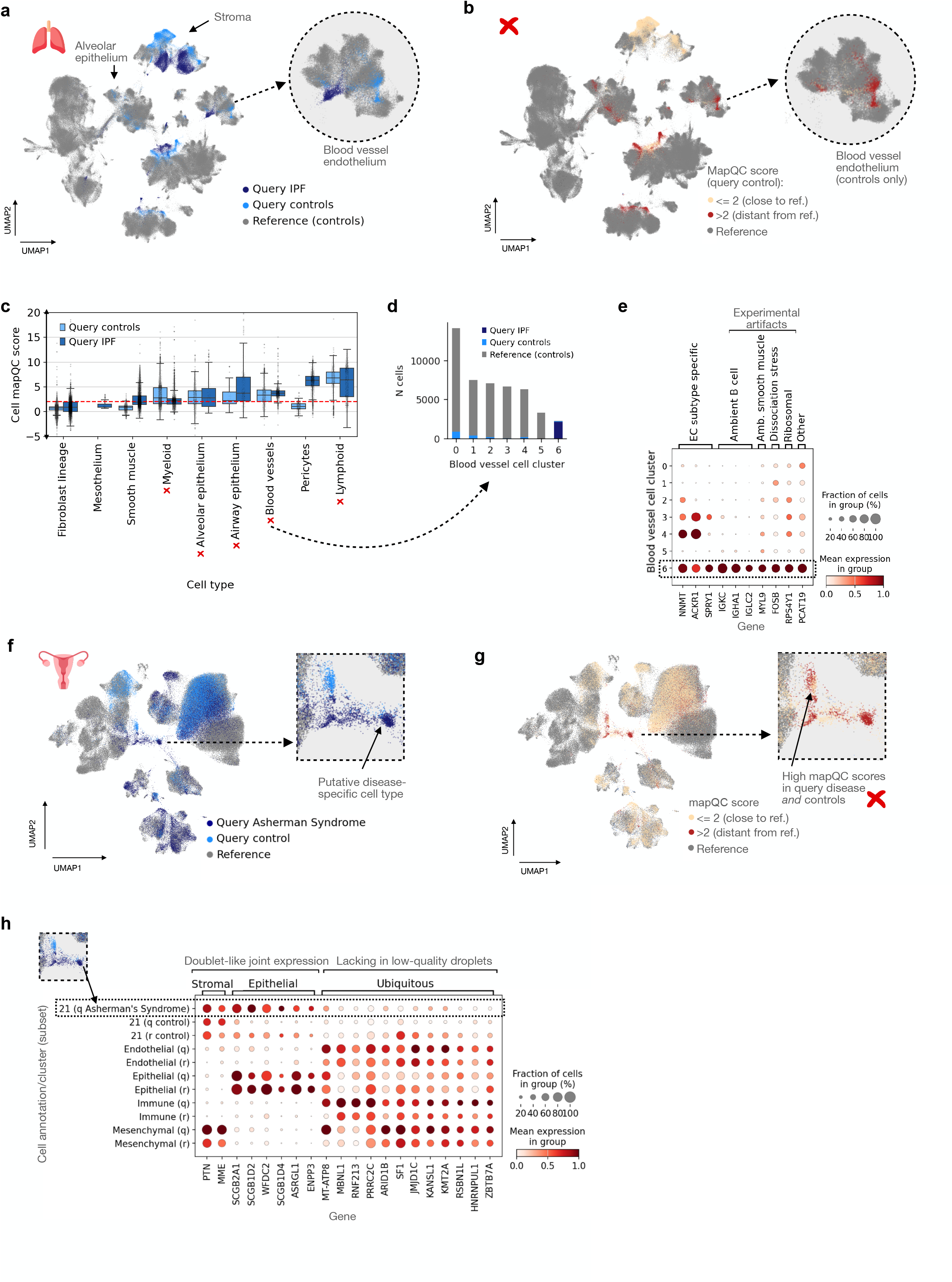
MapQC correctly identifies failed mappings with remaining batch-effect. **a,** UMAP of a query dataset including samples from patients with IPF (dark blue) and controls (light blue) mapped onto the Human Lung Cell Atlas (“reference”, grey). Blood vessel endothelium cells are shown in magnified version in dashed circle. **b,** As (a), with query cells from control samples now colored by mapQC scores, split into low (<2, beige) and high (>2, red) scores. Inset: query control endothelial cells that appear to be mixing with the reference based on the UMAP visualization are in fact distant from the reference, as shown by high mapQC scores. Cells from query IPF samples are not shown. **c,** Query cell mapQC scores shown per cell type. A horizontal red dotted line indicates the cutoff value of 2, above which cells are considered to map distant to the reference. Cell types for which more than half of the cells mapped far from the reference (score>2) are indicated with red crosses. Boxes are included for all groups of at least 100 cells, and show the median and interquartile range. Whiskers extend to the lowest and highest non-outlier points, with data points more than 1.5 times the interquartile range outside the low and high quartile considered outliers. Outliers above 20 or below −5 are not shown for figure legibility, but are shown in Extended Data fig. 2b. **d**, Blood vessel cell cluster composition, broken down by data subset and lung condition (reference, query control, or query IPF). **e**, Expression of genes related to experimental artifacts across blood vessel cell clusters of the joint HLCA reference and query. **f**, UMAP of a query dataset including samples from patients with Asherman syndrome (dark blue) and healthy controls (light blue) mapped onto the Human Endometrial Cell Atlas (HECA “reference”, grey). Cells identified as a disease-specific cell state in the original publication are shown in magnified version in dashed square. **g,** As (f), with query cells now colored by mapQC scores, split into low (<2, beige), and high (>2, red) scores. **h**, Expression of several marker/gene groups among cells in the HECA reference and mapped query, showing marker expression for high-mapQC-score-cluster 21 (split into reference, query control, and query disease), and all other cells split by lineage (and further split into reference “r” or query “q”). For both (e) and (h), expression values were normalized per gene, such that the group with the highest expression was set to 1, and the group with the lowest expression to 0.

To determine whether a large distance from the reference (mapQC score*>*2) in the query control indeed indicates remaining batch effect in the embedding, we selected a subset of cells to analyze in more detail. Specifically, we chose a cell type for which the large distance of the control cells from the reference, as inferred by mapQC, could not be seen in the UMAP. Blood vessel endothelial cells from query control samples accordingly mixed with the reference in the UMAP despite high mapQC scores, whereas query disease cells separated from the reference (**fig. 2a, b**). Mimicking a standard analysis workflow, we clustered the endothelial cells to see if disease cells formed a separate cluster. Indeed, clustering not only resulted in separate clusters for different endothelial cell subtypes, but also in an additional cluster containing almost all query disease endothelial cells (**fig. 2d, Extended Data fig. 2c, d**). Normally, differences between this disease-specific cluster and other clusters would be interpreted as disease-specific expression patterns. However, the top differentially expressed genes, aside from cell type marker genes, consisted of expression changes related to experimental artifacts, including ambient RNA (such as from B cells, *IGKC* and *IGHA1*), dissociation-stress related (*FOSB*), and ribosomal protein (often related with batch effects[6], *RPS4Y1*) genes (**fig. 2e, Extended Data fig. 2e, f**). MapQC thus correctly identifies mapped cells from this cell type as containing remaining batch effects by assigning high mapQC scores to the control cells. It thereby prevents the downstream interpretation of batch effects as disease-specific effects.

As a second use case, we mapped a dataset to the Human Endometrial Cell Atlas (HECA)[17] (**fig. 2f, Extended Data fig. 3a, b**). The query dataset included data from 6 healthy donors and 9 donors with Asherman’s syndrome (AS), a disease of the uterus that can cause infertility[18]. Most query control cells mapped close to the reference (mapQC score*<*2 for 93.9% of cells), suggesting a largely successful removal of batch effects during the mapping (**fig. 2g, Extended Data fig. 3c**). While no clear difference in scores between control and disease cells could be observed at the cell type level, partitioning the cells into clusters did result in the isolation of groups distant from the reference: two clusters (cluster 21 and 25) consistently showed mapQC scores above 2, and consisted largely of cells from the query (94 and 97% of cells, **Extended Data fig. 3c-f**). Interestingly, one of these two clusters (cluster 21) contained 74% of cells from the cell type described as disease-specific “AS epithelium” in the original publication of the query dataset (**fig. 2f, g,** zoom-in), possibly indicating these cells mapped separate from the reference due to disease-specific gene expression patterns.

However, both clusters also contained many cells from query control samples with high mapQC scores, suggesting that the observed distance between these clusters and the reference was at least partly batch-rather than disease-driven (**Extended Data fig. 3d, f**).

To investigate if the query-specific clusters were indeed batch-effect-dominated, we identified cluster-specific differentially expressed genes for cluster 21, containing the putative disease-specific AS epithelium. Up-regulated genes in this cluster revealed a disease-specific gene expression pattern, with the query disease but not query control cells in the cluster showing doublet-like expression of both stroma- and epithelium-specific genes (**fig. 3h**). However, down-regulated genes in the cluster were lowly expressed in both control and disease query cells, while highly expressed across the rest of the cells in the reference and query, showing that differential expression patterns in this cluster were only partly disease-specific (**fig. 3h**). Moreover, cells in both cluster 21 and 25 showed a low number of total counts per cell, did not express marker genes for any specific cell-type (as based on HECA markers) beyond general stromal markers, and showed low *MALAT1* expression in the case of cluster 25 (**Extended Data fig. 3g-i**). Noisy gene expression patterns, which can result in lack of specific marker expression, low total counts, and low *MALAT1* expression are known signs of low-quality “cells”, such as empty droplets[19–21]. Similarly, the co-expression of genes from distinct lineages (stroma, epithelium) observed in the query disease cells is characteristic of doublets, as further supported by the lack of genes uniquely expressed in the cluster. Overall, the two clusters with high mapQC scores appear to be dataset-specific low-quality cells rather than disease-specific cellular phenotypes, as claimed in the original publication. The cell types have thus been correctly flagged by mapQC as a mapping artifact that cannot be used for downstream analysis. In contrast, the remainder of control cells in the query dataset largely show low mapQC scores, suggesting downstream analysis can be performed after removal of the low-quality cells. Thus, mapQC aids in identifying integration failures during the mapping, preventing mis-interpretation of the data.

**Figure 3.**
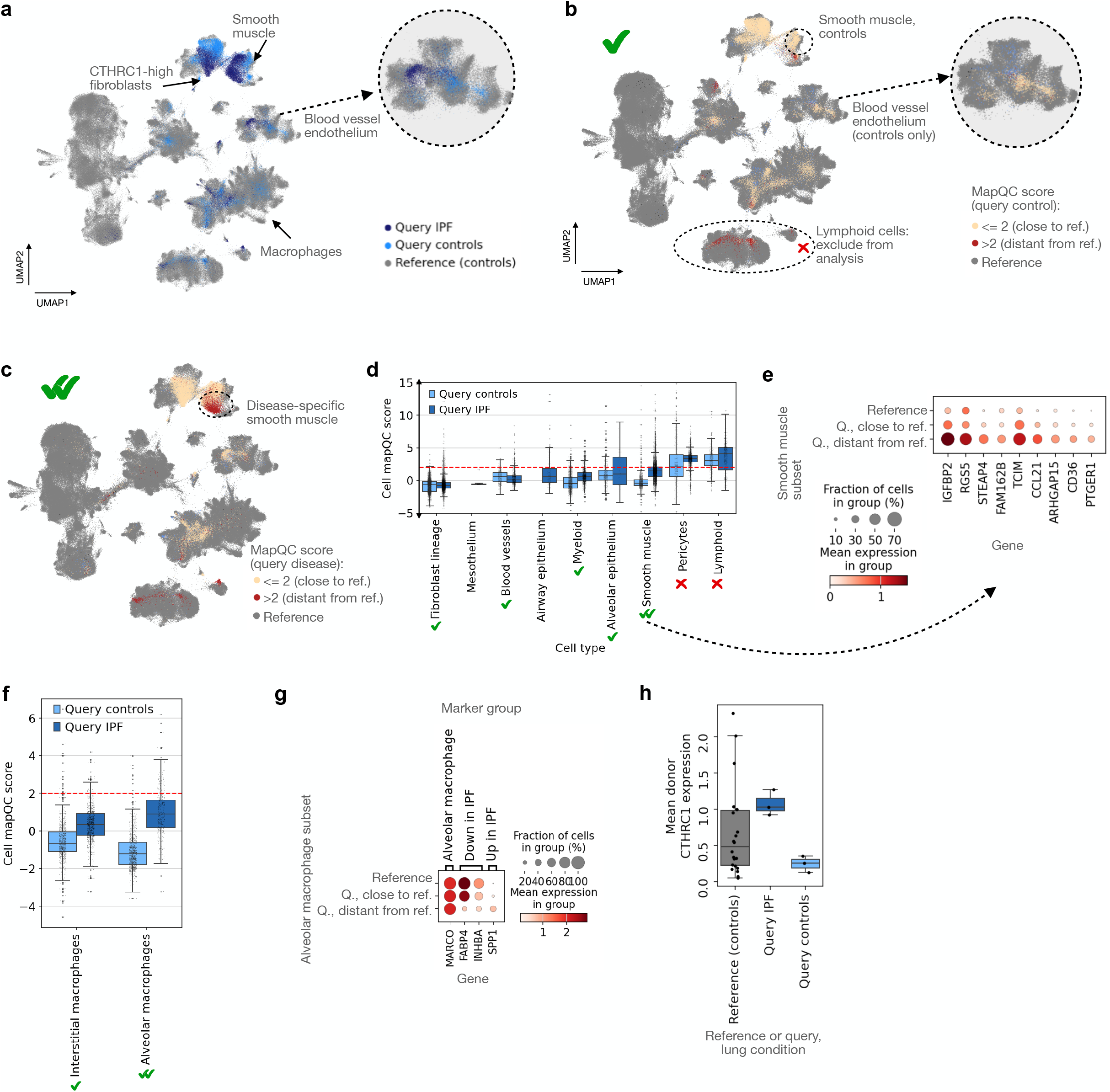
Successful mappings identified by mapQC enable the identification of disease-specific cell states. **a**, UMAP of a query dataset (see fig. 2a) remapped with different parameter settings, including cells from healthy controls from HLCA reference (grey), from query controls (light blue) and query donors with IPF (dark blue). A zoom-in of the UMAP of the blood vessel endothelium is shown. **b**, As (a), with query control cells now colored by mapQC scores, split into low (<2, beige) and high (>2, red) scores. Query disease cells are not shown. **c,** As (b), but now showing query disease cells from IPF patients, while excluding query control cells. **d**, Query mapQC scores from (b) and (c) shown per cell type, split into control and case (IPF) cells. Cell types with more than half of the control cells mapping close to the reference (score <2) are indicated with green ticks, and those with more than half mapping far from the reference (>2) with red crosses. Double green ticks indicate cell types with control cells close to the reference, and disease cells far from the reference. A horizontal red dotted line indicates the cutoff value of 2, above which cells are considered distant from the reference. Boxes are shown for groups with at least 100 cells. Outliers above 15 and below −5 (29 and 64 cells, respectively) are not shown for legibility, but are shown in extended data fig. 4b. **e**, Expression of genes differentially highly expressed among smooth muscle cells distant from the reference (mapQC score >2), shown for smooth muscle cells from the reference and from the query (“Q.”), the latter split into high and low mapQC score groups. **f**, MapQC scores among subgroups of macrophages in the query. **g**, Expression of known alveolar macrophage markers, either general or IPF-specific, among reference alveolar macrophages, and query alveolar macrophages mapped close to or distant from the reference according to mapQC. **h**, Expression of CTHRC1, reported as IPF specific in the query data publication, among cells in the CTHRC1-high fibroblast cluster of the joint reference and query. Mean expression per donor is shown, with only donors with at least 10 cells in the cluster included in the figure. Donors are grouped based on coming from the reference or query, and based on lung condition. For all box plots, the boxes show the median and interquartile range. Whiskers extend to the lowest and highest non-outlier points, with data points more than 1.5 times the interquartile range outside the low and high quartile considered outliers.

### 2.3 Successful mappings enable the identification of disease-specific cell states

To investigate whether mapQC could correctly identify a successful mapping, we remapped the previously discussed query dataset of IPF onto the HLCA, while changing several mapping parameters to non-default settings aiming to improve the mapping (**Methods**). Changing the maximum KL weight parameter only worsened mapping quality (percentage of cells distant from the reference (mapQC score*>*2), in query control samples in original mapping: 28.4%, versus 100% for maximum KL weight 2, 49.1% for maximum KL weight 0.1). In contrast, mapping with a different batch covariate (sample instead of dataset) substantially improved the mapQC scores of the control samples (17.1% of cells distant from the reference). Additionally recovering counts from 69 mapping input genes that had gotten lost during genome annotation harmonization further improved mapping scores (9.3% of cells distant from the reference, **fig. 3a-d, Extended Data fig. 4a, b**, query controls). Lymphoid (NK, B, and T) cells, low in cell number, and pericytes mapped distant from the reference (high mapQC scores) in both control and disease samples, suggesting these cell types still included batch-specific variation in the embedding and should be analyzed with caution. Interestingly, integration quality of NK and T cells was already reported to be poor in the reference itself[6], stressing the importance of a high-quality reference for high-quality mapping. Overall, cells from disease samples were more distant from the reference than those from the controls for several cell types (9.3% versus 23.7% of cells with mapQC score*>*2), indicating disease-specific effects were likely well-preserved during mapping (**fig. 3b-d**, query IPF). Unlike in the previous mapping with query control ECs mapping distant from the reference (**fig. 2a-e**), in this mapping ECs from the query disease samples mapped close to the reference and did not form a separate cluster (**Extended Data fig. 4c**). The previously described batch-specific ambient RNA thus seems to have correctly been ignored during this mapping.

In contrast to the endothelial cells, query smooth muscle cells did map distant from the reference, with 26% of cells from IPF lungs but only 2% of cells from control samples (**fig. 3d**) showing high mapQC scores. Moreover, this result is consistent across study subjects (**Extended Data fig. 4d**). Comparing gene expression between query smooth muscle cells mapped close to or distant from the reference, we found many genes specifically expressed in cells distant from the reference (**fig. 3e**), suggesting a disease-specific smooth muscle phenotype. Indeed, most of these genes also show increased expression in IPF smooth muscle in several other datasets [22–24], confirming the disease state identified by mapQC is not caused by batch-related artifacts in the data (**Extended Data fig. 4e**). Several of these genes, including *IGFBP2, FAM162B*, [25] and *CCL21* [26–28], have been previously associated with lung fibrosis and IPF.

Notably, alveolar macrophages highly expressing *SPP1*, a previously described IPF-specific cell state[6, 24, 29], are also correctly identified as disease-specific based on their distance from the reference, while alveolar macrophages low in *SPP1* and high in *FABP4* and *INHBA* expression, as previously observed in healthy[30, 31], map close to the reference (**fig. 3f,g**). Importantly, in the UMAP of the joint reference and query, macrophages from IPF patients do not visibly map separately from their healthy counterparts (**fig. 3a, b**), confirming that a more quantitative approach - not based on two-dimensional data visualization-is needed to identify cells distant from the reference. Interestingly, query fibroblasts highly expressing *CTHRC1*, reported as disease-specific in the original publication of the mapped dataset[16], map close to the reference according to mapQC, thus not being highlighted as disease-specific. Indeed, inspection of *CTHRC1* expression of samples clustering with *CTHRC1*-high cells from IPF donors shows that 7 healthy donors show a mean *CTHRC1* expression equal to or higher than the IPF donors in the cluster (**fig. 3h, Extended Data fig. 4f-h**). None of these are control samples from the query dataset itself, showing the relevance of mapping to a larger control population (reference atlas) for a more accurate identification of disease-specific phenotypes. Overall, mapQC aids in identifying disease-specific cell populations in a successful query-to-reference mapping.

### 2.4 MapQC outperforms integration metrics

Currently, several metrics are being used to evaluate query-to-reference mappings. These metrics are largely pre-existing integration metrics that were re-purposed as mapping metrics. We here focus specifically on metrics that do not require cell-type labels of the query, that allow for cell-level scores, and that can be used in combination with any mapping method.

First, a number of metrics aim to quantify preservation of the query manifold structure before and after mapping. For example, Azimuth’s cluster preservation score compares clustering of the query before and after mapping[12]. Similarly, Azimuth’s mapping score[12] and Symphony’s within-query k-NN-correlation[9] (“wiq-kNN-corr”) compare query cells’ nearest query neighbors before and after mapping. Whereas these metrics could be used to highlight loss of information after mapping, they do not evaluate the quality of the integration of the query and the reference. This is confirmed by these metrics’ evaluation of the best and worst mapping, according to mapQC, of the previously discussed IPF dataset to the HLCA. Whereas for the worst mapping the query control cells separate completely from the reference even in the UMAP, scores for these metrics (cluster preservation score, wiq-kNN-corr) are nearly indistinguishable from those for the best mapping (**fig. 4a-c**). Similarly, donor LISI scores, which aim to quantify the mixing of query donors in the embedding after mapping, show no clear differences between the two mappings. In contrast, mapQC identifies 100% of query control cells as distant from the reference in the bad mapping.

**Figure 4.**
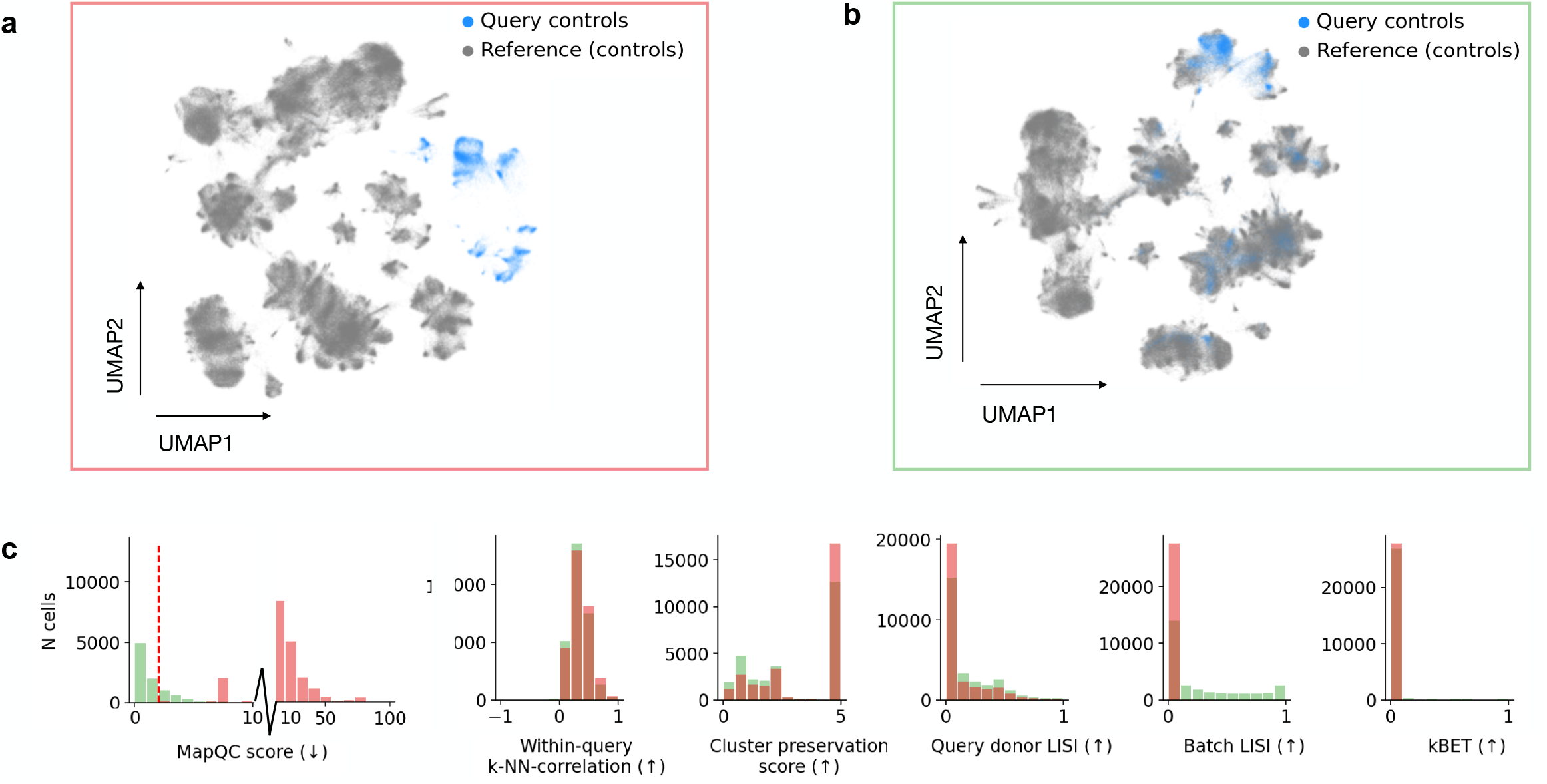
MapQC is the only metric that correctly distinguishes a failed from a successful mapping, compared to commonly used integration and query-to-reference mapping metrics. **a**, UMAP of a failed; and **b**, a successful query-to-reference mapping. Only cells from the HLCA reference (grey) and query healthy controls (light blue) were included for this benchmark. **c**, Scores of mapQC and 5 other commonly used integration and mapping metrics. Scores shown in red were calculated calculated on the mapping shown in (a), scores in green on the mapping shown in (b). A vertical red dotted line is shown for the cutoff-value of mapQC scores separating good from bad mappings. Arrows in the x-axis labels show whether a lower or a higher value indicates good mapping for the specific metric. For mapQC scores, the x-axis is split into 0-10 and 10-100, to enable easy visual comparison of the scores for the two different mappings (c and d, left panel).

A fourth metric, kBET[32], which quantifies the preservation of batch composition in neighborhoods across the embedding, shows the lowest possible score for almost all cells in both embeddings. kBET assumes that query and reference proportions should be constant across larger regions in the embedding, an assumption that is not expected to hold given a reference highly diverse in sample types (different anatomical regions sampled, different protocols used, etc.), making the metric overly sensitive (**fig. 4**).

The final metric, the LISI score, while showing a clearer difference between the two mappings, is highly sensitive to the abundance of different batches. The value of the score is determined by the number of draws from a cell’s nearest neighbors until a “batch” (query donor for “donor LISI”, or reference or query for “batch LISI”) is observed twice. Indeed, “batch LISI”, the LISI score that is widely used to compare batch integration performance across different integrations methods [13], shows dramatically different scores when down-sampling query cells to 1% or even 10% of the original number of cells, whereas mapQC scores generally stay stable (**Extended Data fig. 5**).

Unlike for mapQC, a clear interpretation of the absolute values of any of these metrics is lacking, making a non-arbitrary and generalizable cutoff value for bad integration as well as a successful integration impossible. MapQC is thus the first evaluation metric that can be readily interpreted and is suited for a quality-check of any given mapping to a large-scale reference, reliably identifying both successful and failed mappings.

Crucially, all the discussed metrics ignore natural variation between samples, making their integration scores less accurate and often ill-founded: these metrics do not distinguish between normal versus abnormal inter-sample variation, but qualify any lack of inter-batch mixing as bad. In contrast, mapQC is the first metric to incorporate a natural inter-sample variation estimate into its score calculation, thus enabling a more precise distinction between failed and successful mappings.

## 3 Discussion

Large-scale reference atlases are being constructed at an increasingly fast pace, with atlases including a large set of healthy or unperturbed controls now available for several organs, model systems, and cell types in both human and mouse [1, 33–37]. This development ushers in a transformed way of analyzing single-cell data: rather than analyzing datasets in isolation, they will now routinely be analyzed in the context of large-scale references[1, 7]. Indeed, reference atlases are already being used widely for the annotation and interpretation of new data [2–4]. Correct usage of reference atlases is therefore more important than ever: incorrect mapping of a new query dataset to an existing reference will lead to misinterpretation of the query. In this study we presented mapQC, a metric to evaluate query-to-reference mappings in an automated and interpretable fashion. Leveraging the availability of a large-scale reference, mapQC takes into account normal inter-sample variation by constructing local null-distributions of inter-sample distances. This results in the first biologically grounded and interpretable distance metric to evaluate query-to-reference mappings. We show that mapQC correctly identifies remaining batch effects in query-to-reference mappings based on large distances of query controls to the reference, and does so better than standard integration metrics. MapQC moreover correctly identifies successful mappings, enabling the discovery of true disease-specific cell states. It thus facilitates a more reliable usage of reference atlases for the analysis of new data.

We note that the use of mapQC is bound to several limitations. First, it requires the availability of control samples in the query dataset that were generated with a protocol similar to the case samples. Whereas the requirement of control samples in a dataset might incur extra costs, including controls is considered good scientific practice in any scientific experiment and is currently also routinely done in single-cell experiments[38– 40]. Moreover, it has been shown previously that the availability of a limited set of matched controls, even when mapping to a reference atlas of more controls, significantly reduces the chance of false positives in disease state identification[5]. Second, mapQC’s assumption that the reference covers most of the variation in the control population is only likely to hold for large-scale reference atlases. Therefore, mapQC is less suitable for evaluating the mapping of one dataset to a single other dataset. Its ability to accurately distinguish batch-effects from query-specific cell states depends on the number and diversity of donors in the reference. For example, if the reference only covers a subset of the healthy population (e.g. only people of European descent) and the query contains data from a different population, mapQCs assumptions are violated, causing it to conflate biology with batch effects. Similarly, the quality of the reference can affect mapQC’s performance: if inter-sample distances in the reference are driven by batch effects, mapQC will not be able to identify remaining batch effects of the same strength in the query after the mapping. Finally, mapQC makes the assumption that the extent of batch effects is similar in the query control and query disease samples. While this assumption has a clear rationale (sample collection and processing, which are a major source of batch effect, are often similar across samples from the same dataset), it might not always hold. However, a perfect confounding between experimental conditions and covariates of interest (e.g. disease status) cannot be solved by any data-driven metric, and would require the use of prior knowledge for correct analysis of the data.

One important feature of mapQC is its calculation of inter-sample distance null distributions at the neighborhood level. This neighborhood-level approach accounts for cell-state specific levels of inter-sample variation. Indeed, we observed substantial differences in inter-sample distance between and even within cell types (**Extended Data fig. 1**). This variation might have multiple sources. First, it could be caused by biological phenomena, such as cell type or state-specific expression quantitative trait loci (eQTLs) that cause different levels of inter-individual variaton, or cell type-specific sensitivity to changes in environment (e.g. disease, aging))[6, 41, 42]. Second, it might have computational sources, such as a better representation of cell-type specific gene expression variance by the embedding model for more abundant cell types. In both cases, taking this variation into account is essential when quantifying a query cell’s distance to the reference.

For our mapping of IPF data to the HLCA reference, we showed an example of improving the quality of a mapping by tuning several mapping parameters and by increasing the quality of the query input data. However, in many other mapping cases parameter tuning did not lead to a sufficiently good mapping due to remaining batch effects in the data. Moreover, tuning possibilities are limited for all methods[8–10]. We therefore identify a need for new query-to-reference methods that allow for more tuning flexibility and a stronger batch-correction of the query or even the reference data when mapping.

Whereas we only showed the use of mapQC in the context of query-to-reference mapping of a query to a large-scale healthy reference, there are several other possible uses of mapQC that have not been show-cased here. For example, mapQC could be used during the construction of large-scale reference atlases themselves, to compare the integration of specific datasets to the rest of the datasets in the reference. Similarly, mapQC can be employed in the context of smaller references (e.g. one large reference dataset instead of an integrated atlas). In all these cases, it is important to keep in mind - and validate where possible-the assumptions that mapQC is built on (e.g. that the subjects from the query dataset are a sample from a healthy population that was also sampled in the reference).

In this study, we introduced mapQC, a metric designed to evaluate the quality of query-to-reference mappings in single-cell analysis. MapQC is the first metric tailored specifically to the query-to-reference mapping scenario, leveraging information about inter-sample variation present in the reference to accurately evaluate a query’s integration into the reference. We demonstrated that existing integration metrics fail to distinguish failed from successful mappings. In contrast, mapQC correctly detects failed mappings, and accurately evaluates successful ones, enabling the discovery of true disease-associated cell states. The emergence of large-scale single-cell atlases has fundamentally shifted the way single-cell data analysis is performed, with reference mapping now being a key first step in annotation and interpretation of new data. The need for a rigorous evaluation of such mappings has thus become more critical than ever. We anticipate that mapQC will serve as an essential tool to ensure the reliability of reference-based analyses, advancing our ability to identify disease-specific cell states and ultimately advancing our understanding of health and its changes in disease.

## 4 Methods

### 4.1 Selecting neighborhoods

Inter-sample distances are calculated at the level of neighborhoods. As centers of the neighborhoods, a subset of query cells are selected, for which the k nearest neighbors are calculated. The query center cells are randomly selected per cluster of the joint embedding of the reference and query, with the number of query cells sampled per cluster being proportional to the fraction of query and the fraction of reference cells in each cluster. This approach ensures that both cell types abundant in the reference, but less abundant in the query and *vice versa* are well sampled. Specifically, the fraction of the total number of sampled cells per cluster *c* is calculated as follows:

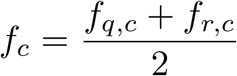

where *f_q,c_* and *f_r,c_* are the fraction of total query and total reference cells, respectively, that are part of cluster *c*. In cases where the target number of query cells to sample from a cluster is lower than the total number of query cells in that cluster, the remainder of cells is sampled from the remaining clusters proportional to their sampling fractions *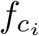*. In total, 500 neighborhoods are selected per query dataset. This number can be adapted based on the size of the joint reference and query. Note that as the sizes of the neighborhoods are large (see “setting the neighborhood size k” below), these neighborhoods are expected to overlap in many cases. A neighborhood-based approach is also taken by a related method, Milo[43], which aims to quantify local differential abundance between (groups of) samples, e.g. of different conditions. However, whereas they are similar in their neighborhood-based approach, mapQC and Milo have different goals: Milo quantifies changes in cell abundance, while mapQC is deliberately made insensitive to abundance (number of cells per sample), as these can change depending on sample processing protocol, anatomical location sampled, single-cell method, etc., which are independent of the quality of a specific mapping. Rather, mapQC quantifies the local *distance* between samples, which reflects the mixing of the samples’ cells in the embedding space, as will be discussed below.

### 4.2 Setting the neighborhood size k

For all organ reference atlases analyzed in this paper, the number of neighbors k per neighborhood was set to 500. The size of k should be set such that each neighborhood encompasses something between a cell type and a cell substate. Setting the k too large will result in inter-sample distances being overshadowed by inter-cell type distances, in cases where cell type sizes are smaller than k, which is not what mapQC tries to quantify. Moreover, a large k is more likely to result in a bi-or multi-modal distribution of cells for a single sample in the embedding space of the neighborhood, which defies mapQC’s assumptions for inter-sample distance calculations. In contrast, setting k too small will fail to capture the general mixing behavior of reference samples for a given cellular phenotype. Note that the k can be adapted to the type of atlas. For example, for single-cell-type atlases (e.g. a NK cell atlas), we recommend using a much larger neighborhood size, such that the chosen k covers biologically relevant subgroups of cells. Additionally, the k of individual neighborhoods can be adapted if these neighborhoods do not pass specific filters, as further described below.

### 4.3 Sample and neighborhood filtering and adapting individual neighborhood sizes

Briefly, the following filtering and parameter change steps are followed before proceeding to inter-sample distance: 1) in each neighborhood, remove samples with too few cells; 2) identify neighborhoods with too few reference samples; 3) increase neighborhood size for neighborhoods from (2); 4) remove neighborhoods with too few reference samples even after increasing the neighborhood size. More specifically, for step 1, in each neighborhood, samples with fewer than 10 cells are excluded from distance calculations. For steps 2 and 3, for neighborhoods with fewer than 3 reference samples that passed filtering, the k is increased, as the baseline distance between reference samples cannot be reliably assessed for these neighborhoods with the standard neighborhood size. The cells from poorly mapped query datasets will map far from the reference, resulting in such query-only neighborhoods. For these neighborhoods, the final k is set to 1.1 times the neighborhood size needed to reach the minimum number of reference samples per neighborhood. To limit computation time and to prevent neighborhoods from covering multiple cell types at the same time, the maximum k is set to 10 times the original k (e.g. 5000 for a default k of 500). For step 4, a neighborhood is excluded from downstream calculations if it still does not include sufficient reference samples at the maximum k. Note that in many cases, at least part of the cells in those neighborhoods will still be covered by other neighborhoods.

### 4.4 Calculating a sample’s distance to the reference

To calculate a sample’s distance to the reference in a given neighborhood *h*, the procedure as described below is followed. First, samples that do not pass the above-described filtering are excluded from calculations, such that *S_h_* is the set of all samples in the neighborhood *h* that passed filtering. Then, for each sample *s_i_* ∈ *S_h_*, its distance to each sample *s_j_* ∈ *S_h,r_* is calculated, where *i ≠ j*, and with *S_h,r_* being the set of samples in *S_h_* coming from the reference. To calculate the distance between samples, several distance metrics were explored. First, earth-mover’s distance or “Wasserstein distance” was tried, a distance metric which is often used to calculate the distance between two distributions, such as two sets of cells[44–46]. However, calculating this distance is computationally expensive, and since the number of times this distance has to be calculated per mapping is approximately

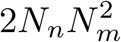

where *N_n_* is the number of neighborhoods and *N_m_* is the mean number of samples per neighborhood, using this distance was not computationally feasible: it took more than 48 hours to calculate for a single mapping. As a second alternative, a simple mean pairwise distance metric was considered. This metric can be used to quantify the distance between two sets of cells, but also depends on the variance within these sets: the distance between two groups of cells is zero if and only if their means are equal and they both have a variance of zero. As the primary goal of the metric was to quantify distance between distributions (i.e. lack of mixing) rather than (changes in) variance, we decided against using a simple pairwise distance. Finally, energy distance or “E-distance”[14] was investigated as a metric to quantify distances between samples. E-distance is a statistical measure of the distance between distributions, intuitively reflecting the signal-to-noise ratio of a specific effect (e.g. of a perturbation) in a given set of data points[47]. E-distance has been used previously to detect effects of specific perturbations[15, 48]. Briefly, the E-distance is calculated by taking the mean Euclidean distance between two groups of cells, and then subtracting the mean distance within the two groups. As this metric seemed to be a more accurate formalization of our aim (identifying shifts in distributions between reference and query), we decided to use E-distance as our default distance metric, although other distance metrics are also available to the MapQC user. Specifically, the E-distance between *s_i_* and *s_j_ E*(*s_i_, s_j_*) in a given neighborhood is calculated, considering only the cells in that neighborhood, as:

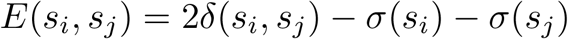

where *δ*(*s_i_, s_j_*) is the mean pairwise distance between the cells from sample *s_i_* and sample *s_j_*:

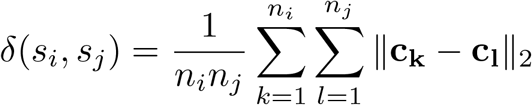

and *σ*(*s_i_*) is the mean pairwise distance between cells from sample *s_i_*:

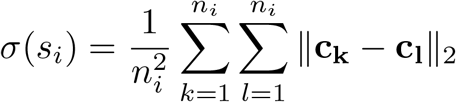

and where *n_i_* is the number of cells in *s_i_*. As distances between pairs of cells, the euclidean distance ∥**c_k_** − **c_l_**∥**_2_** between the cells in the embedding space is used.

The distance of a given query sample *s_i,q_* to the reference is then calculated as the mean distance of that sample to all samples from the reference:

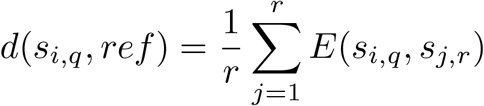

where *r* is the number of samples from the reference in neighborhood *h* that have passed filtering |*S_h,r_*|, and *s_j,r_* is a sample from the reference. Importantly, to calculate the distance of query samples to the reference, no distances between samples from the same dataset are ever used during the calculation (as the query by definition comes from a different dataset than the reference). To not bias distance to the reference for reference samples, which could be smaller within dataset than between datasets, we therefore exclude reference samples from the same dataset when calculating the distance of any reference sample to the reference, such that:

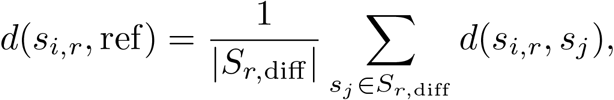

where *S*_*r*,diff_ is the set of reference samples in *S_h_* that are from a different dataset than that of *s_i_*.

### 4.5 Normalizing distances across neighborhoods

Inter-sample distances can differ considerably from neighborhood to neighborhood, both as a result of differences in cell density across the embedding space, and due to lower or higher levels of mixing between samples in different regions of the embedding, e.g. due to cell types showing more or less inter-individual heterogeneity. The goal of mapQC is to establish whether or not query samples mix “normally” with the reference, which is approximated by checking if they are at a “normal” distance from the reference. Here, the reference itself is considered the baseline and is therefore used to define what normal inter-sample distances are. Thus, distances to the reference are normalized or “Z-scored” as follows, with all variables below being specific to the neighborhood under consideration:

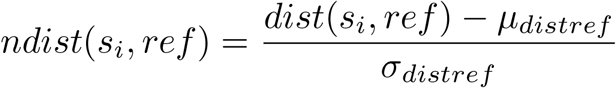

Where

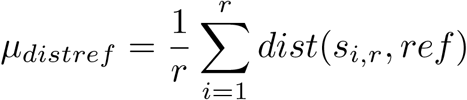

And

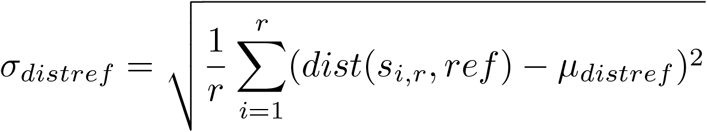

such that the normalized distances of the reference samples to the reference in any given neighborhood have a mean of 0 and a standard deviation of one. In cases of extreme outliers among the reference samples, these outliers are removed from the above calculation of the mean and standard deviation, such that normalized distances are not heavily affected by single outliers in the data. Outliers are defined as any points more than 3.5 times the inter-quartile range (IQR) above the third quartile of reference sample distances to the reference, or more than 3.5 times the IQR below the first quartile.

### 4.6 MapQC score calculation

The mapQC score for a given query cell *c_i,s_* quantifies how well the sample *s* from which that cell came mixes with the reference in the region around cell *c_i,s_*. The mapQC score is therefore calculated as the mean distance to the reference of sample *s* across the set of all neighborhoods that cell *c_i,s_* is part of (*N_c_*):

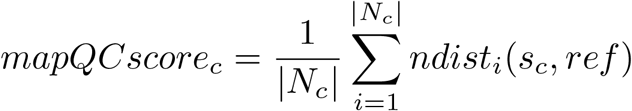

Note that due to the stochastic nature of the mapQC algorithm (sampling a random subset of query cells as neighborhood centers), in some cases cells will not have a mapQC score. This can be e.g. because all neighborhoods it was part of were filtered out, or because it was not part of any neighborhood. If this is the case for many cells, the neighborhood size or number might need to be increased.

### 4.7 Query-to-reference mapping with scArches

As currently available large healthy reference atlases are almost all integrated with scVI or related methods[1, 17, 34, 49–51], all reference atlases used in this paper had also been integrated with scVI[11] or scANVI[52]. Mapping was therefore done exclusively with scArches[8], currently the only query-to-reference mapping method available that is compatible with conditional variational autoencoder-based integrations such as those generated with scVI and scANVI. For the mapping, raw counts were used, subsetted to the input genes of the base integration model. For the mapping of the IPF lung query dataset[16], raw counts were taken directly from the Human Lung Cell Atlas[6]. For re-mapping of that same data with counts recovered from 69 genes that had gone missing during gene name conversion for upload of the HLCA to cellxgene, the original counts were used as made available with the original query publication (GSE132771). For the mapping of the Asherman’s syndrom query dataset[18], controls that were included in the data but that were generated as part of a different dataset originally were excluded, as mapQC assumes similar sample handling across samples from the same dataset. As base integration models, the reference model of the HLCA[6] was used for the lung IPF data query, and the scANVI reference model of the HECA developed for reference mapping [17] was used for the endometrial Asherman Syndrome data query. Batch covariate was set as indicated in the main text, as dataset or sample for the IPF query, and as sample for the Asherman Syndrome query. No cell type labels were included for the mapping. The following parameter settings were used for the mapping: number of epochs: 500, weight decay: 0, and maximum KL weight set to default 1 unless specified otherwise.

### 4.8 Count pre-processing, clustering, and visualization of joint reference and query

For the HLCA core reference and the mapped query, normalized and log-transformed counts were directly taken from the HLCA[6]. For the HECA, non-control samples in the reference (i.e. from donors with endometriosis) were excluded from all downstream analyses, as mapQC works under the assumption that the reference includes control samples only. For both the HECA reference and the Asherman Syndrom query dataset, raw counts were normalized to 10,000 counts per cell, and then log-transformed using the natural log and a pseudo-count of 1. For all mapping-derived joint reference and query embeddings, a knn-graph was constructed based on the embedding using the scanpy package[53] setting the number of neighbors k=30 for the HECA, and k=50 for the HLCA mappings, and with default settings otherwise. To cluster the data, leiden clustering[54] was performed based on the kNN-graph, using the scanpy package with default settings. For data visualization, a UMAP embedding[55] was calculated based on the kNN graph using the scanpy package with default settings.

### 4.9 Cell type label transfer from reference to query

To annotate cells in the query, cell type label transfer was performed from the reference to the query for all mapped datasets. For each cell in the query, its k (30 for HECA, 50 for HLCA) nearest neighbors in the reference were calculated based on the joint embedding derived from the mapping. To reduce computation time, the pynndescent package[56] (specifically the PyNNDescentTransformer function) was used to identify, and calculate Euclidean distances to the k nearest neighbors. For each query cell, the probability of the cell being of cell type *l* (*p_l_*) was then calculated as follows:

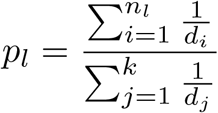

where *k* is the total number of neighbors, *n_l_* is the number of neighbors with cell type label l, and *d_i_* is the distance of neighbor i to the query cell. Each cell was then labeled as the cell type with the highest probability for that cell.

### 4.10 Downstream analysis of IPF query-to-reference mappings

For the first mapping (used as an example of failed mapping) of the IPF query dataset[16], with default mapping parameters and treating all samples as a single batch (batch covariate “dataset”), endothelial cells were clustered as follows: first, all endothelial cells (excluding lymphatic endothelium) were selected based on their level 2 annotation in the reference (“blood vessels”) or based on the transferred annotation in the query. The resulting subset of cells was re-clustered by first re-calculating a kNN-graph based on the embedding as described above, with the number of neighbors k=30, and then performing Leiden clustering as described above with resolution=0.3. To identify differentially expressed genes, log-normalized gene expression of the cluster containing almost exclusively and almost all endothelial cells from the donors with IPF (cluster 6) was compared to gene expression in all other endothelial cells using a t-test (**Supplementary table 1**). P-value correction for multiple tests was done using the Benjamini-Hochberg procedure. The resulting gene list was loosely filtered based on the following criteria, to select the most relevant differentially expressed genes: minimum fraction of cells within the cluster expressing the gene: 0.25, minimum fold change: 1 (i.e. only up-regulated genes), maximum fraction of cells outside of the cluster expressing the gene: 0.5. Finally, to ensure that differentially expressed genes identified as possible batch-effect related genes were actually used during the mapping and could therefore have changed the embedding of the cells, we subsetted the resulting gene list to only genes that were used during the mapping. For visualization (**fig. 2e, Extended Data fig. 2e,f**), the top 30 genes that passed filtering were taken, after which genes not used for mapping were excluded, such that 10 genes remained. Marker genes shown in the figure were taken from the HLCA marker gene set[6]. For the second mapping (used as an example of a successful mapping), with the batch covariate set to sample and a larger set of genes included for the mapping, endothelial cells were clustered exactly as described for the previous mapping. For the smooth muscle analysis, query smooth muscle cells with high mapQC scores (*>*2) were compared to query smooth muscle cells with low mapQC scores (*<*2) (**Supplementary table 2**), after which the results were filtered as described above (minimum in-group fraction 0.25, maximum out-group fraction 0.5) and keeping only genes with Benjamini-Hochberg multiple testing-corrected p-values below 0.05. From the resulting list of 50 genes, the genes most unique for the high mapQC score query cells (compared to all other stromal subgroups) were selected based on visual inspection and used for visualization (**fig. 3e**) and for comparison to other datasets (**Extended Data fig. 4e**). For comparison to other datasets, all datasets in the HLCA[6] that included smooth muscle cells from both healthy donors and donors with IPF were taken (“Sheppard 2020” is the mapped query[16], other studies: Banovich 2020[22, 23], Kaminski 2020[24], Misharin Budinger 2018[29]), and pre-processed gene counts from the published HLCA were used for visualization of gene expression (**Extended Data fig. 4e**). For the analysis of *CTHRC1*-high cells, all cells labeled as stromal except for mesothelium cells were re-clustered in the same way as described above for the endothelial cells. To determine which samples from both the reference and query showed high *CTHRC1* expression, cells were then subset to only the fibroblast cluster containing the far majority of *CTHRC1*-high cells. *CTHRC1*-expression for cells in this cluster is shown per donor in **fig. 3h** and **Extended Data fig. 4h**.

### 4.11 Downstream analysis of Asherman Syndrome query-to-reference mapping

To compare gene expression of cells in cluster 21 (containing many query cells with high mapQC scores) to that of other cells, a differential gene expression analysis was performed comparing cells in cluster 21 to all other cells in the joint reference and query, including only genes that were detected in at least one cell in both the reference and the query, using a t-test (**Supplementary table 3**). P-value correction for multiple tests was done using the Benjamini-Hochberg procedure. The results were then loosely filtered such that only genes expressed in at least 25% of cells within cluster 21, and expressed in at most 50% of cells outside of cluster 21, and with a mean expression higher inside than outside the cluster were kept (i.e. only up-regulated genes). The resulting set of 12 genes was largely widely expressed in either the stromal or the epithelial subset of all cells, while being jointly expressed in cluster 21. The 8 genes for which this was most clearly the case are shown in **fig. 2h** and labeled as “stromal (up in 21)” and “epithelial (up in 21 q. disease)”. A second differential gene expression analysis was performed to identify genes down-regulated in query cells from cluster 21 compared to query cells only from all other clusters as described above (**Supplementary table 4**). The resulting list of differentially expressed genes was stringently filtered, such that only genes being expressed in at least 50% of cells outside of the cluster, and in no more than 50% of cells within the cluster were kept, and only genes with a mean expression higher outside of the cluster than inside. Of the resulting 19 genes, 7 were excluded as these were expressed in subsets of cells outside of the cluster based on visual inspection. The remaining 12 genes are shown in **fig. 2h** and labeled as “ubiquitous (down in cl. 21 query)”.

### 4.12 Comparison to commonly used integration and mapping metrics

To compare mapQC to existing mapping and integration metrics, we calculated scores for five other metrics: within-query-kNN correlation, cluster preservation, donor LISI, batch LISI, and kBET. Importantly, in order to make a fair comparison to mapQC, in cases where the metric as previously used consisted of a single value per dataset, we used the metric at the cell level, i.e. before summarizing per-cell values in a single dataset score. Finally, we used parameter settings as used in the original metric calculations to obtain the best possible version of the score, resulting in slightly different pre-processing per metric as further specified below. As an example of a good mapping, we took the best-performing mapping of the IPF query dataset[16] in terms of mapQC scores, i.e. with sample as batch covariate, maximum KL divergence set to 1 during mapping, and with 69 more input genes recovered as compared to the cellxgene HLCA counts matrix. As an example of a bad mapping, we took the worst-performing mapping of the same dataset, i.e. with dataset as batch covariate, the maximum KL divergence set to 2 for the mapping, and using the HLCA cellxgene counts matrix (i.e. 69 input genes missing).

The **within-query kNN-correlation** (wiq-kNN-corr) score, used to quantify mapping quality with the query-to-reference method Symphony, was calculated as previously described[9]. This score aims to quantify preservation of the local embedding structure of the query by comparing the distance to each cell’s k nearest neighbors in two embeddings. Specifically, a principal component analysis (PCA) was performed on the normalized, log-transformed query count matrix. The first 50 principal components were subsequently used to calculate Euclidean distances to the 500 nearest neighbors for each query cell. Then, the same distances (between all query cells and their original 500 nearest neighbors) were calculated in the embedding space resulting from the mapping. To determine how well the neighborhood structure was preserved for each cell after mapping, the distances in the two different embeddings were compared using Spearman correlation, such that perfect correlation would result in the maximum score of 1 for a given cell.

Similar to the wiq-kNN-corr score, the **cluster preservation** score aims to quantify embedding similarity in two different embeddings of the same data using clustering. The cluster preservation score is a mapping quality metric used for Seurat’s Azimuth query to reference mapping method[10, 12]. Specifically, the normalized, logtransformed gene expression matrix of the query dataset was scaled per gene such that each gene’s mean and variance were 0 and 1, respectively. Then, a PCA was performed on the scaled count matrix, and the first 50 components were used for the rest of the analysis (embedding 1). The 20 nearest neighbors of each cell were calculated based on the PCA embedding, and cells were then clustered using the Louvain algorithm[57] with the resolution parameter set to 0.6. To calculate the cluster entropy in the original PCA embedding, for each cell the frequency of each cluster label was determined among its 20 nearest neighbors. Cluster entropy *H* for this cell was then calculated using Shannon entropy:

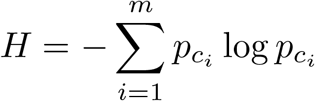

where m is the number of distinct cluster labels observed among the cells nearest neighbors, and *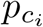* is the proportion of cells among the nearest neighbors from cluster *c_i_*.

Then, the same was done using the mapping-based embedding (embedding 2), calculating each cell’s 20 nearest neighbors and calculating cluster entropy for each cell based on its neighbors and the original cluster labels (from embedding 1). To compare how well the new embedding had preserved the structure of the original embedding, the difference in clustering entropy was calculated for each cell as:

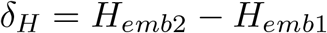

The final cluster preservation score per cell was then calculated by setting the score to the maximum value of 5 if the entropy was lower in the new than in the original embedding (i.e. *δ_H_ <* 0), or re-scaling otherwise:

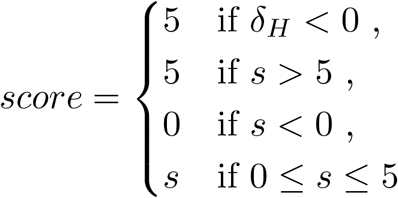

Where

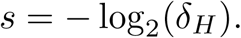

The **kBET** score was calculated as previously described[13]. Specifically, the 50 nearest neighbors were calculated for each query cell, after which a chi-square test was used to compare the observed reference and query cell frequency in the cell’s neighborhood to the overall reference and query cell frequency across all cells. The resulting p-values were then reported as cell-level scores.

The Local Inverse Simpson’s Index (LISI) scores, **donor LISI** and **batch LISI**, were calculated as previously described[9, 58]. LISI quantifies how many cells can be drawn from a group of neighbors before the same group (e.g. donor, batch) is observed twice. Specifically, for donor LISI scores, a kNN graph was constructed with number of neighbors k=30 using the embedding from the mapping, and taking only the query cells into account. The donor LISI score per cell was then calculated (ranging from 1 to N, where N is the number of donors in the query), after which the results were rescaled such that they ranged from 0 to 1, as follows:

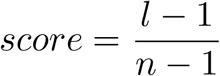

where l is the original, non-normalized LISI score, and n is the total number of groups (donors in this case).

Similarly, the batch LISI score was calculated on the combined query and reference, first constructing a kNN graph with number of neighbors k=90 using the mapping-based embedding, after which the LISI score was calculated for each cell, now using only two groups: reference or query. The final batch LISI score per cell was then obtained by normalizing for the donor LISI score.

For determining the robustness of the batch LISI score and mapQC score to changes in query cell numbers, the “successful” mapping of the mapped IPF dataset[16] was used, i.e. the mapping as shown in **fig. 3**, and only query control cells were included and randomly down-sampled twice to both 10% and 1% of the total number of query cells.

### 4.13 Software versions

For all preprocessing, MapQC runs, and downstream analyses: anndata: 0.10.9; leidenalg: 0.10.1; pynndescent: 0.5.10; python: 3.9.0; scanpy: 1.10.3; scikit-learn: 1.3.0; scipy: 1.11.2; umap-learn: 0.5.3. For all query-to-reference mappings, and for the comparison to existing metrics: python: 3.8, scanpy: 1.9.2; scarches 0.5.7, scib-metrics: 0.5.1, scvi-tools 0.20.1.

## Supplementary information

Supplementary Information includes supplementary table 1 to 4.

## Acknowledgments

We thank Lennard Halle, Michaela Muüller and Chelsea Bright for feedback on the manuscript, Dominik Klein for help with implementing Jax-based Sinkhorn distance as part of mapQC’s distance metric comparisons, Eljas Roellin for the suggestion to use Energy distance as a distance metric, and Ronan Le Gleut from the Core Facility Statistical Consulting at Helmholtz Munich for statistical advice.

## Code availability

The mapQC package can be found under https://github.com/theislab/mapqc/. All code used for results in this manuscript can be found under https://github.com/theislab/mapqc_reproducibility/tree/v0.1.0-manuscript.

## Authors’ contributions

L.S. and F.J.T. conceived of the initial idea for the project. L.S., M.L. and D.S. conceptualized the project. L.S. implemented the project, with contributions from M.L. and D.S. A.A.M. implemented and organized code for comparison to existing integration metrics. J.E. contributed to reference atlas search and setup. L.S. wrote the manuscript with contributions from A.A.M., D.S, and F.J.T. All authors read and provided feedback on the manuscript.

---

**Extended Data Figure 1.**
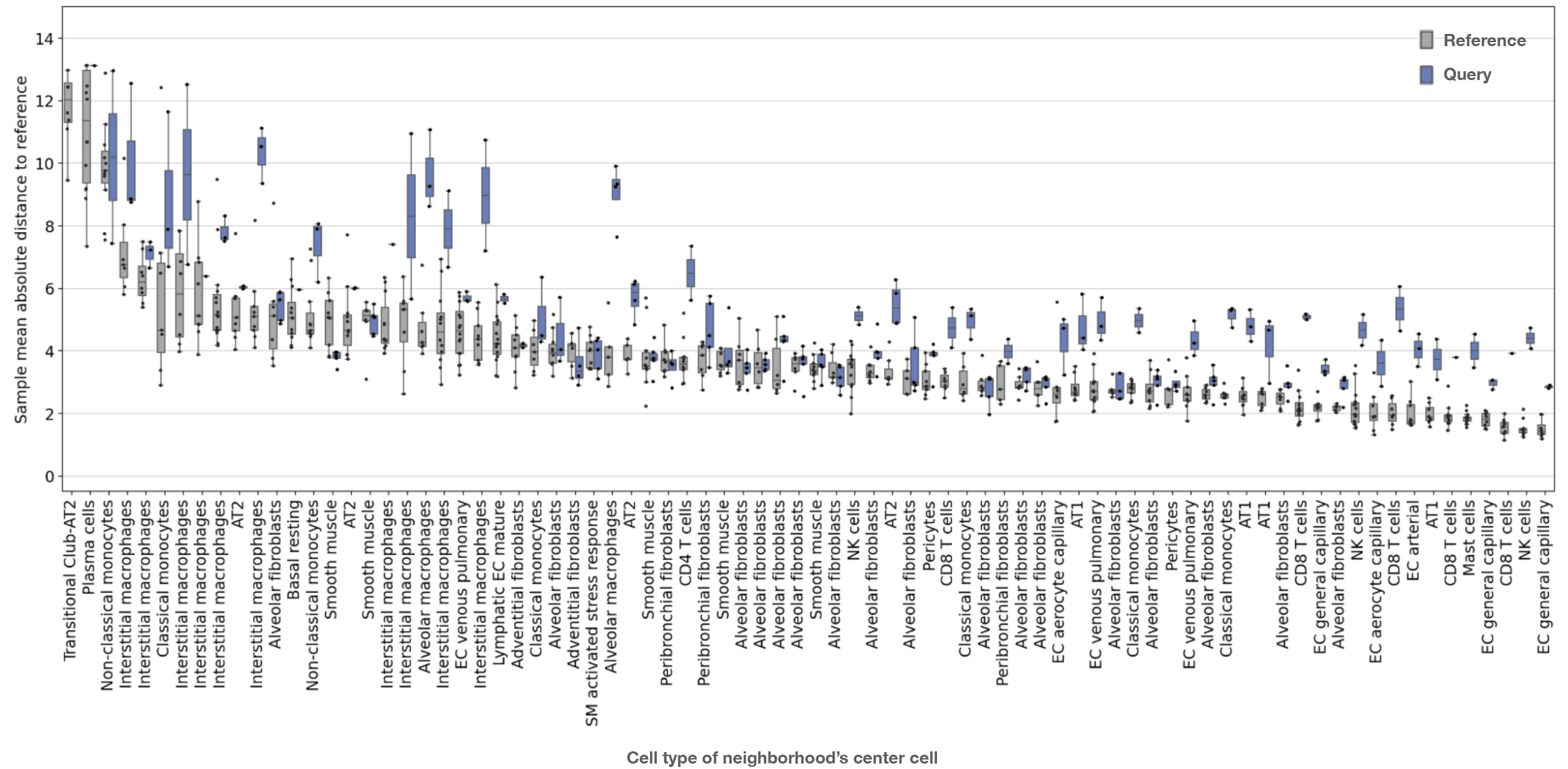
Sample absolute distances to the reference highly vary across neighborhoods. Each neighborhood is labeled based on the cell type of its center cell. Distances are shown for reference samples (grey) and query samples (blue). Neighborhoods are ordered based on the mean distance to the reference among reference samples. Boxes show the median and interquartile range. Whiskers extend to the lowest and highest non-outlier points, with data points more than 1.5 times the interquartile range outside the low and high quartile considered outliers.

**Extended Data Figure 2.**
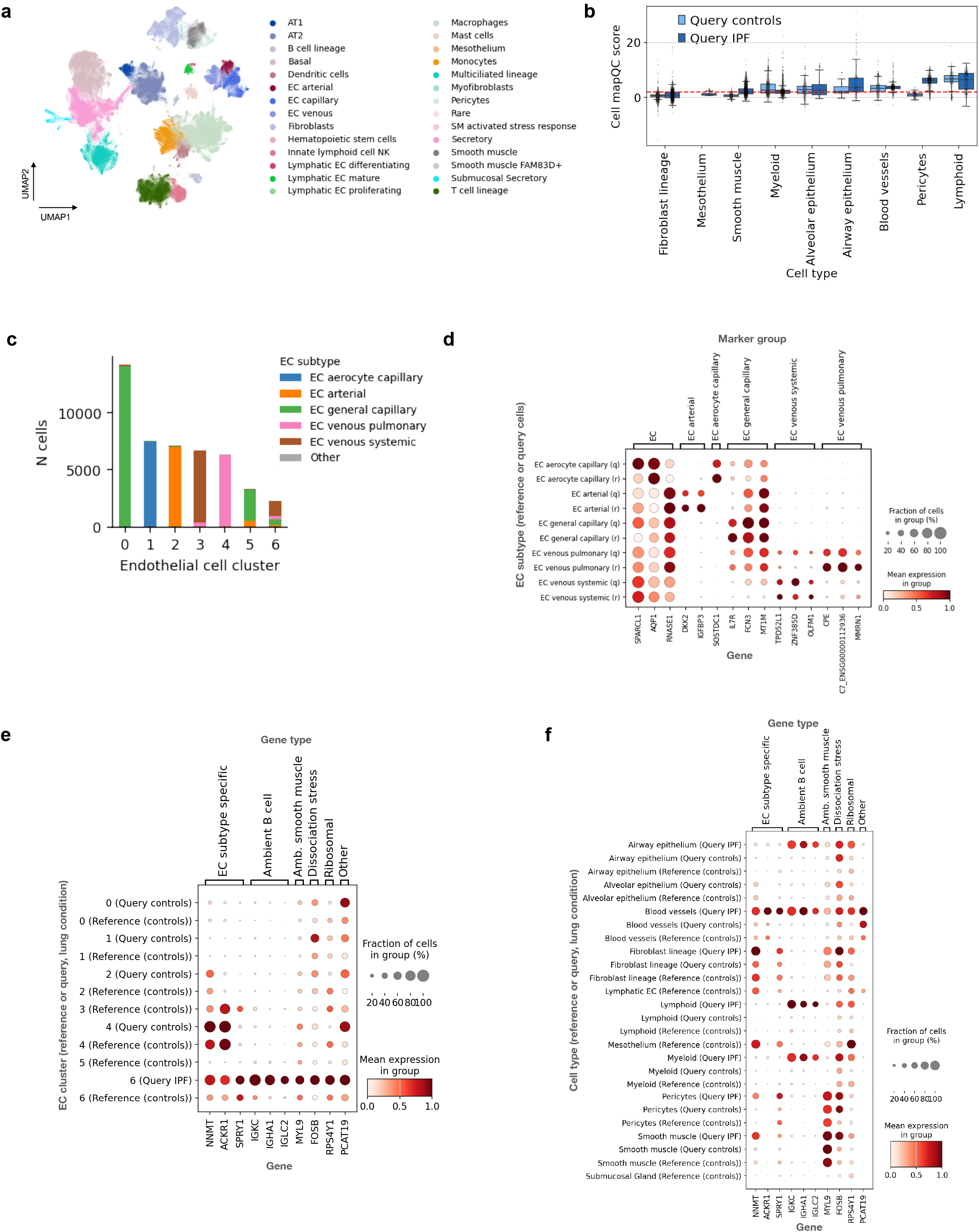
Clustering and marker gene expression among query and reference cell subsets of a failed mapping. **a,** UMAP of a query dataset with data from IPF donors and controls mapped onto the Human Lung Cell Atlas reference. Cells are colored by cell type. Query data was annotated by performing label transfer from the reference to the query. **b,** Query cell mapQC scores shown per cell type. A horizontal red dotted line indicates the cutoff value of 2, above which cells are considered not to mix well with the reference. Boxes are included for all groups of at least 100 cells, show the median and interquartile range. Whiskers extend to the lowest and highest non-outlier points, with data points more than 1.5 times the interquartile range outside the low and high quartile considered outliers. **c,** Cluster composition by EC subtype of blood vessel endothelial cell clusters, **d,** Expression of blood vessel subtype marker genes per annotation, split by reference (annotation) and query. Cell annotations with fewer than 50 cells were excluded. **e,** Expression of genes differentially highly expressed in cluster 6-dominated by query IPF cells. Expression is shown for all EC clusters, split by reference (all controls), query controls and query IPF. Groups with fewer than 50 cells were excluded. **f,** Expression across all cell types of genes differentially expressed in query IPF blood vessel cells as compared to other blood vessel cells (see (d)). Note that query stromal cells (pericytes, fibroblasts, smooth muscle and mesothelium) largely came from separate, stromal lineage-enriched samples, therefore showing independent ambient RNA patterns. Only groups with at least 200 cells are shown. For (c-e), gene expression values were normalized per gene, such that minimum and maximum expression were set to 0 and 1, respectively.

**Extended Data Figure 3.**
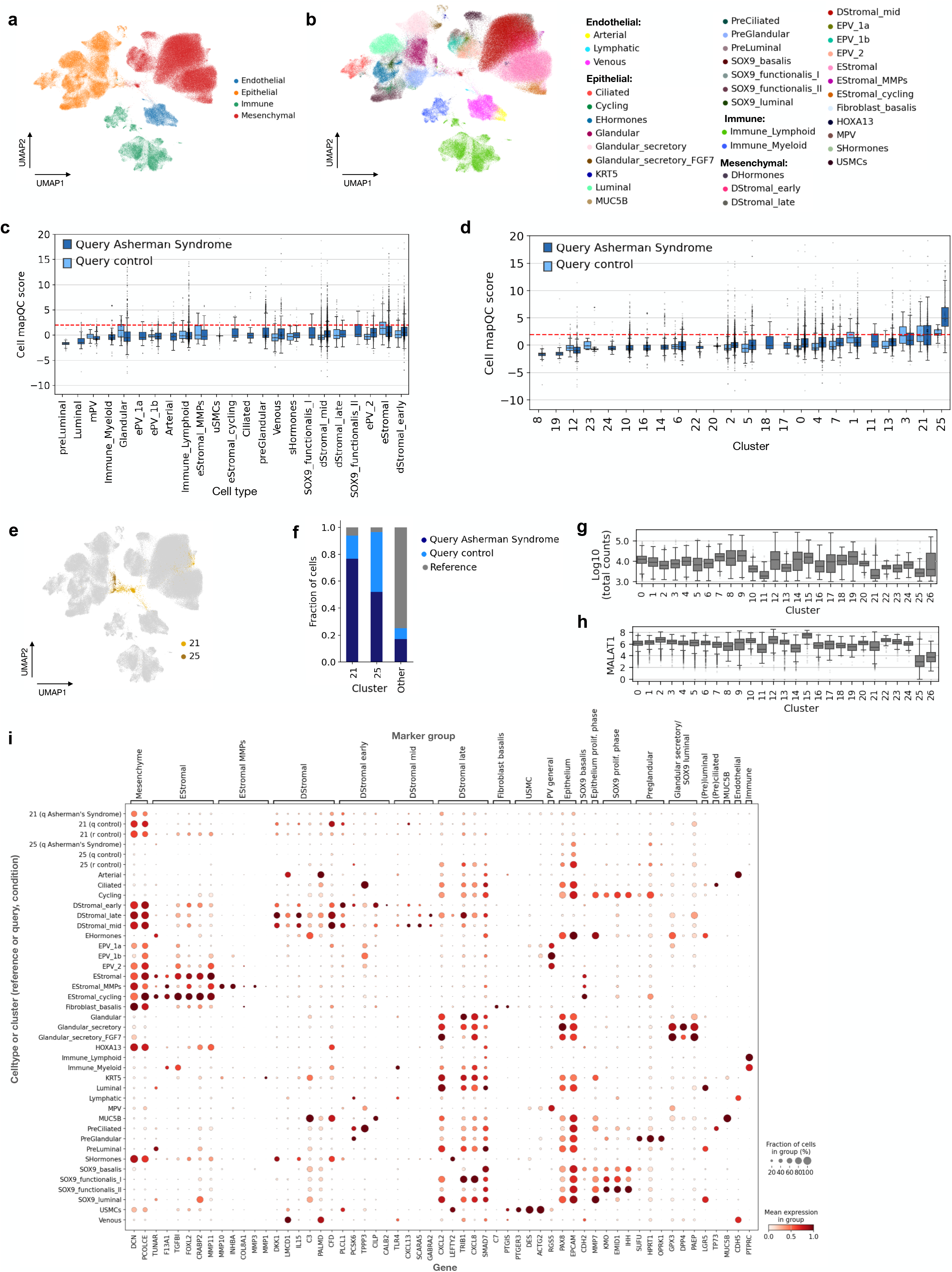
Details of the mapping of a query dataset of donors with Asherman syndrome onto the Human Endometrial Cell Atlas (HECA) reference. **a**, UMAP of the joint reference and query. Cells are colored by lineage. Query cells were annotated based on label transfer from the reference. **b**, As (a), now coloring cells by cell type. **c**, MapQC scores of mapped query cells, grouped by cell type. Cells are split into cells from control donors (light blue) or cells from donors with Asherman Syndrome (dark blue). A red dashed horizontal line shows the mapQC score cutoff value of two. **d**, As (c), now grouping cells by cluster. **e**, UMAP of the joint reference and query (as in (a)), now showing the location of the two clusters with the highest mapQC scores (21 and 25, yellow and ochre yellow, respectively) in the UMAP. **f,** Composition of cluster 21, 25, and the rest of the cells by group, with groups being reference (grey), query controls (light blue), and query Asherman syndrome (dark blue). **g**, Log 10 of the total number of RNA counts per cell in the query dataset, grouping cells by cluster, showing only clusters with at least 50 cells from the query data. **h,** Expression of MALAT1 per cell in the query datasets, grouping cells by cluster as in (g). **i**, Marker gene expression (marker gene set based on HECA markers) among all cell types in the joint reference and query, showing cells from cluster 21 and 25 separately, and splitting the latter two into cells from the reference (“r control”), from query control donors (“q control”), and query donors with Asherman syndrome (“q Asherman’s syndrome). Expression values are normalized per gene, such that the minimum and maximum expression are set to 0 and 1, respectively. For all box plots, boxes are included for all groups with at least 50 cells. The boxes show the median and interquartile range. Whiskers extend to the lowest and highest non-outlier points, with data points more than 1.5 times the interquartile range outside the low and high quartile considered outliers.

**Extended Data Figure 4.**
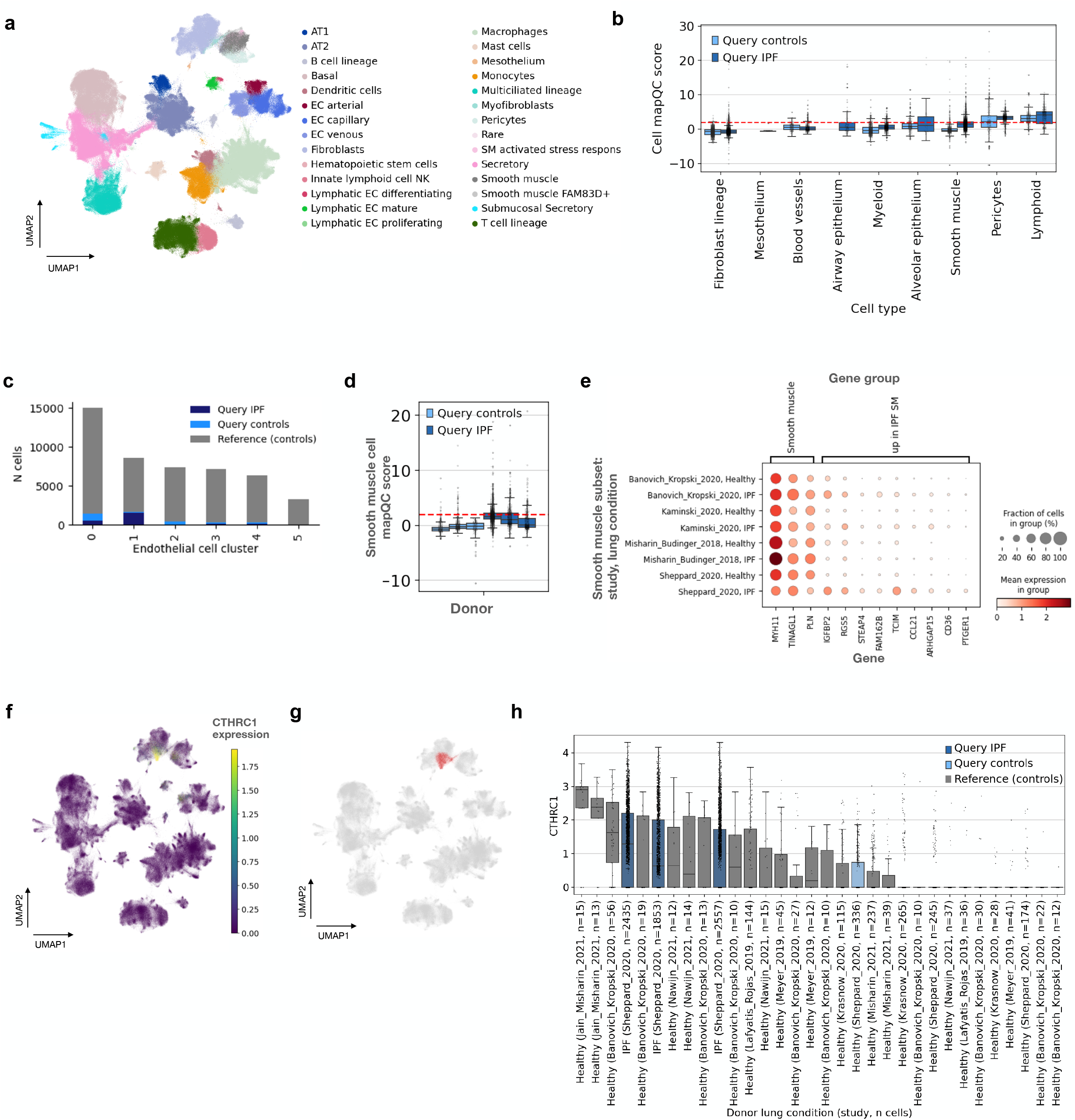
Details of a successful mapping of a query dataset of donors with IPF onto the HLCA. **a**, UMAP of the joint reference and query, colored by cell type. Query cells were annotated by performing label transfer from the query to the reference. **b**, Query mapQC scores of mapped cells shown per cell type, split into control and case (IPF) cells. A horizontal red dotted line indicates the cutoff value of 2, above which cells are considered not to mix well with the reference. Boxes are shown for groups with at least 100 cells. **c**, Composition of blood vessel endothelial cell clusters by subgroup, breaking cells down by reference or query and lung condition (reference controls (grey), query controls (light blue), and query IPF (dark blue)). **d**, MapQC scores among smooth muscle cells, splitting cells by donor, with boxes for query controls colored in light blue (left three boxes) and for query donors with IPF in dark blue (right three boxes). **e**, Expression of general smooth muscle cell markers and genes up-regulated in high mapQC score smooth muscle cells from the mapped query dataset (“Sheppard_2020”). Expression is shown across several datasets from studies including donors with IPF and controls. Cells are grouped by study, and then by lung condition. **f**, UMAP of the joint reference and query, colored by *CTHRC1* expression, a marker identified as IPF-specific in the original query publication. **g** UMAP as in (e), now showing the location of the fibroblast cluster high in CTHRC1. **h**, CTHRC1 expression among donors with at least 10 cells in the cluster from (f). Boxes are grey for donors from the reference, light blue for query control donors, and dark blue for query donors with IPF. X tick labels show the donor’s lung condition, the study from which the donor came, and the number of cells the donor had in the cluster. For box plots in (c) and (g), the boxes show the median and interquartile range. Whiskers extend to the lowest and highest non-outlier points, with data points more than 1.5 times the interquartile range outside the low and high quartile considered outliers.

**Extended Data Figure 5.**
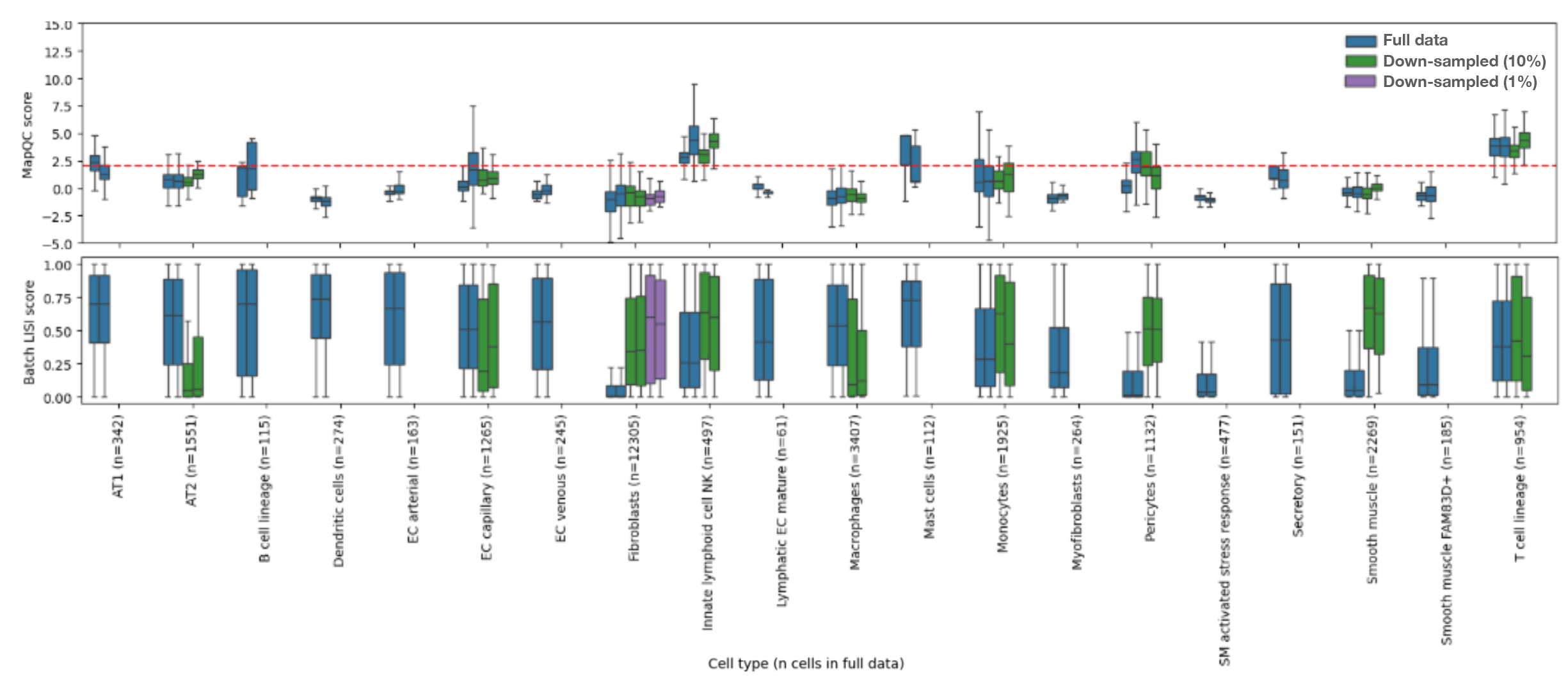
The number of query cells dramatically changes the batch LISI scores, while having a small effect on mapQC scores. Scores were calculated twice on the full query control data (blue boxes), on the control data randomly down-sampled twice to 10% of cells (green boxes), and to 1% of cells (purple boxes). Results are show only for cell types that had at least 50 cells remaining after a specific down-sampling. Top panel: mapQC scores, with a dashed line showing the threshold value of 2. Bottom: batch LISI scores, treating reference and query as batches. Boxes show the median and interquartile range. Whiskers extend to the lowest and highest non-outlier points, with data points more than 1.5 times the interquartile range outside the low and high quartile considered outliers, which are not shown.

## References

[1] Hrovatin, K., Sikkema, L., Shitov, V.A., Heimberg, G., Shulman, M., Oliver, A.J., Mueller, M.F., Ibarra, I.L., Wang, H., Ramírez-Suástegui, C., He, P., Schaar, A.C., Teichmann, S.A., Theis, F.J., Luecken, M.D.: Considerations for building and using integrated single-cell atlases. Nature Methods 22(1), 41–57 (2024) 10.1038/s41592-024-02532-y

[2] Galimberti, M., Nucera, M.R., Bocchi, V.D., Conforti, P., Vezzoli, E., Cereda, M., Maffezzini, C., Iennaco, R., Scolz, A., Falqui, A., Cordiglieri, C., Cremona, M., Espuny-Camacho, I., Faedo, A., Felsenfeld, D.P., Vogt, T.F., Ranzani, V., Zuccato, C., Besusso, D., Cattaneo, E.: Huntington’s disease cellular phenotypes are rescued non-cell autonomously by healthy cells in mosaic telencephalic organoids. Nature Communications 15(1) (2024) 10.1038/s41467-024-50877-x

[3] Lang, N.J., Gote-Schniering, J., Porras-Gonzalez, D., Yang, L., De Sadeleer, L.J., Jentzsch, R.C., Shitov, V.A., Zhou, S., Ansari, M., Agami, A., Mayr, C.H., Hooshiar Kashani, B., Chen, Y., Heumos, L., Pestoni, J.C., Molnar, E.S., Geeraerts, E., Anquetil, V., Saniere, L., Wögrath, M., Gerckens, M., Lehmann, M., Yildirim, A.O., Hatz, R., Kneidinger, N., Behr, J., Wuyts, W.A., Stoleriu, M.G., Luecken, M.D., Theis, F.J., Burgstaller, G., Schiller, H.B.: Ex vivo tissue perturbations coupled to single-cell rna-seq reveal multilineage cell circuit dynamics in human lung fibrogenesis. Science Translational Medicine 15(725) (2023) 10.1126/scitranslmed.adh0908

[4] Yuan, F., Gasser, G.N., Lemire, E., Montoro, D.T., Jagadeesh, K., Zhang, Y., Duan, Y., Ievlev, V., Wells, K.L., Rotti, P.G., Shahin, W., Winter, M., Rosen, B.H., Evans, I., Cai, Q., Yu, M., Walsh, S.A., Acevedo, M.R., Pandya, D.N., Akurathi, V., Dick, D.W., Wadas, T.J., Joo, N.S., Wine, J.J., Birket, S., Fernandez, C.M., Leung, H.M., Tearney, G.J., Verkman, A.S., Haggie, P.M., Scott, K., Bartels, D., Meyerholz, D.K., Rowe, S.M., Liu, X., Yan, Z., Haber, A.L., Sun, X., Engelhardt, J.F.: Transgenic ferret models define pulmonary ionocyte diversity and function. Nature 621(7980), 857–867 (2023) 10.1038/s41586-023-06549-9

[5] Dann, E., Cujba, A.-M., Oliver, A.J., Meyer, K.B., Teichmann, S.A., Marioni, J.C.: Precise identification of cell states altered in disease using healthy single-cell references. Nature Genetics 55(11), 1998–2008 (2023) 10.1038/s41588-023-01523-7

[6] Sikkema, L., Ramírez-Suástegui, C., Strobl, D.C., Gillett, T.E., Zappia, L., Madissoon, E., Markov, N.S., Zaragosi, L.-E., Ji, Y., Ansari, M., Arguel, M.-J., Apperloo, L., Banchero, M., Bécavin, C., Berg, M., Chichelnitskiy, E., Chung, M.-i., Collin, A., Gay, A.C.A., Gote-Schniering, J., Kashani, B.H., Inecik, K., Jain, M., Kapellos, T.S., Kole, T.M., Leroy, S., Mayr, C.H., Oliver, A.J., Papen, M., Peter, L., Taylor, C.J., Walzthoeni, T., Xu, C., Bui, L.T., Donno, C.D., Dony, L., Faiz, A., Guo, M., Gutierrez, A.J., Heumos, L., Huang, N., Ibarra, I.L., Jackson, N.D., Murthy, P.K.L., Lotfollahi, M., Tabib, T., Talavera-López, C., Travaglini, K.J., Wilbrey-Clark, A., Worlock, K.B., Yoshida, M., Chen, Y., Hagood, J.S., Agami, A., Horvath, P., Lundeberg, J., Marquette, C.-H., Pryhuber, G., Samakovlis, C., Sun, X., Ware, L.B., Zhang, K., Berge, M., Bossé, Y., Desai, T.J., Eickelberg, O., Kaminski, N., Krasnow, M.A., Lafyatis, R., Nikolic, M.Z., Powell, J.E., Rajagopal, J., Rojas, M., Rozenblatt-Rosen, O., Seibold, M.A., Sheppard, D., Shepherd, D.P., Sin, D.D., Timens, W., Tsankov, A.M., Whitsett, J., Xu, Y., Banovich, N.E., Barbry, P., Duong, T.E., Falk, C.S., Meyer, K.B., Kropski, J.A., Pe’er, D., Schiller, H.B., Tata, P.R., Schultze, J.L., Teichmann, S.A., Misharin, A.V., Nawijn, M.C., Luecken, M.D., and, F.J.T.: An integrated cell atlas of the lung in health and disease. Nature Medicine 29(6), 1563–1577 (2023) 10.1038/s41591-023-02327-2

[7] Lotfollahi, M., Hao, Y., Theis, F.J., Satija, R.: The future of rapid and automated single-cell data analysis using reference mapping. Cell 187(10), 2343–2358 (2024) 10.1016/j.cell.2024.03.009

[8] Lotfollahi, M., Naghipourfar, M., Luecken, M.D., Khajavi, M., Büttner, M., Wagenstetter, M., Avsec, Ž., Gayoso, A., Yosef, N., Interlandi, M., Rybakov, S., Misharin, A.V., Theis, F.J.: Mapping single-cell data to reference atlases by transfer learning. Nature Biotechnology 40(1), 121–130 (2021) 10.1038/s41587-021-01001-7

[9] Kang, J.B., Nathan, A., Weinand, K., Zhang, F., Millard, N., Rumker, L., Moody, D.B., Korsunsky, I., Raychaudhuri, S.: Efficient and precise single-cell reference atlas mapping with symphony. Nature Communications 12(1) (2021) 10.1038/s41467-021-25957-x

[10] Hao, Y., Hao, S., Andersen-Nissen, E., Mauck, W.M., Zheng, S., Butler, A., Lee, M.J., Wilk, A.J., Darby, C., Zager, M., Hoffman, P., Stoeckius, M., Papalexi, E., Mimitou, E.P., Jain, J., Srivastava, A., Stuart, T., Fleming, L.M., Yeung, B., Rogers, A.J., McElrath, J.M., Blish, C.A., Gottardo, R., Smibert, P., Satija, R.: Integrated analysis of multimodal single-cell data. Cell 184(13), 3573–358729 (2021) 10.1016/j.cell.2021.04.048

[11] Lopez, R., Regier, J., Cole, M.B., Jordan, M.I., Yosef, N.: Deep generative modeling for single-cell transcriptomics. Nature Methods 15(12), 1053–1058 (2018) 10.1038/s41592-018-0229-2

[12] Azimuth — azimuth.hubmapconsortium.org. https://azimuth.hubmapconsortium.org/. [Accessed 10-12-2024]

[13] Luecken, M.D., Büttner, M., Chaichoompu, K., Danese, A., Interlandi, M., Mueller, M.F., Strobl, D.C., Zappia, L., Dugas, M., Colomé-Tatché, M., Theis, F.J.: Benchmarking atlas-level data integration in single-cell genomics. Nature Methods 19(1), 41–50 (2021) 10.1038/s41592-021-01336-8

[14] Rizzo, M.L., Székely, G.J.: Energy distance. WIREs Computational Statistics 8(1), 27–38 (2015) 10.1002/wics.1375

[15] Replogle, J.M., Saunders, R.A., Pogson, A.N., Hussmann, J.A., Lenail, A., Guna, A., Mascibroda, L., Wagner, E.J., Adelman, K., Lithwick-Yanai, G., Iremadze, N., Oberstrass, F., Lipson, D., Bonnar, J.L., Jost, M., Norman, T.M., Weissman, J.S.: Mapping information-rich genotype-phenotype landscapes with genome-scale perturb-seq. Cell 185(14), 2559–257528 (2022) 10.1016/j.cell.2022.05.013

[16] Tsukui, T., Sun, K.-H., Wetter, J.B., Wilson-Kanamori, J.R., Hazelwood, L.A., Henderson, N.C., Adams, T.S., Schupp, J.C., Poli, S.D., Rosas, I.O., Kaminski, N., Matthay, M.A., Wolters, P.J., Sheppard, D.: Collagen-producing lung cell atlas identifies multiple subsets with distinct localization and relevance to fibrosis. Nature Communications 11(1) (2020) 10.1038/s41467-020-15647-5

[17] Marečková, M., Garcia-Alonso, L., Moullet, M., Lorenzi, V., Petryszak, R., Sancho-Serra, C., Oszlanczi, A., Icoresi Mazzeo, C., Wong, F.C.K., Kelava, I., Hoffman, S., Krassowski, M., Garbutt, K., Gaitskell, K., Yancheva, S., Woon, E.V., Male, V., Granne, I., Hellner, K., Mahbubani, K.T., Saeb-Parsy, K., Lotfollahi, M., Prigmore, E., Southcombe, J., Dragovic, R.A., Becker, C.M., Zondervan, K.T., Vento-Tormo, R.: An integrated single-cell reference atlas of the human endometrium. Nature Genetics (2024) 10.1038/s41588-024-01873-w

[18] Santamaria, X., Roson, B., Perez-Moraga, R., Venkatesan, N., Pardo-Figuerez, M., Gonzalez-Fernandez, J., Llera-Oyola, J., Fernández, E., Moreno, I., Salumets, A., Vankelecom, H., Vilella, F., Simon, C.: Decoding the endometrial niche of asherman’s syndrome at single-cell resolution. Nature Communications 14(1) (2023) 10.1038/s41467-023-41656-1

[19] Ilicic, T., Kim, J.K., Kolodziejczyk, A.A., Bagger, F.O., McCarthy, D.J., Marioni, J.C., Teichmann, S.A.: Classification of low quality cells from single-cell rna-seq data. Genome Biology 17(1) (2016) 10.1186/s13059-016-0888-1

[20] Heumos, L., Schaar, A.C., Lance, C., Litinetskaya, A., Drost, F., Zappia, L., Lücken, M.D., Strobl, D.C., Henao, J., Curion, F., Aliee, H., Ansari, M., Badiai-Mompel, P., Büttner, M., Dann, E., Dimitrov, D., Dony, L., Frishberg, A., He, D., Hediyeh-zadeh, S., Hetzel, L., Ibarra, I.L., Jones, M.G., Lotfollahi, M., Martens, L.D., Müller, C.L., Nitzan, M., Ostner, J., Palla, G., Patro, R., Piran, Z., Ramírez-Suástegui, C., Saez-Rodriguez, J., Sarkar, H., Schubert, B., Sikkema, L., Srivastava, A., Tanevski, J., Virshup, I., Weiler, P., Schiller, H.B., Theis, F.J.: Best practices for single-cell analysis across modalities. Nature Reviews Genetics 24(8), 550–572 (2023) 10.1038/s41576-023-00586-w

[21] Montserrat-Ayuso, T., Esteve-Codina, A.: High content of nuclei-free low-quality cells in reference single-cell atlases: a call for more stringent quality control using nuclear fraction. BMC Genomics 25(1) (2024) 10.1186/s12864-024-11015-5

[22] Habermann, A.C., Gutierrez, A.J., Bui, L.T., Yahn, S.L., Winters, N.I., Calvi, C.L., Peter, L., Chung, M.-I., Taylor, C.J., Jetter, C., Raju, L., Roberson, J., Ding, G., Wood, L., Sucre, J.M.S., Richmond, B.W., Serezani, A.P., McDonnell, W.J., Mallal, S.B., Bacchetta, M.J., Loyd, J.E., Shaver, C.M., Ware, L.B., Bremner, R., Walia, R., Blackwell, T.S., Banovich, N.E., Kropski, J.A.: Single-cell rna sequencing reveals profibrotic roles of distinct epithelial and mesenchymal lineages in pulmonary fibrosis. Science Advances 6(28) (2020) 10.1126/sciadv.aba1972

[23] Natri, H.M., Del Azodi, C.B., Peter, L., Taylor, C.J., Chugh, S., Kendle, R., Chung, M.-i., Flaherty, D.K., Matlock, B.K., Calvi, C.L., Blackwell, T.S., Ware, L.B., Bacchetta, M., Walia, R., Shaver, C.M., Kropski, J.A., McCarthy, D.J., Banovich, N.E.: Cell type-specific and disease-associated eqtl in the human lung (2023) 10.1101/2023.03.17.533161

[24] Adams, T.S., Schupp, J.C., Poli, S., Ayaub, E.A., Neumark, N., Ahangari, F., Chu, S.G., Raby, B.A., DeIuliis, G., Januszyk, M., Duan, Q., Arnett, H.A., Siddiqui, A., Washko, G.R., Homer, R., Yan, X., Rosas, I.O., Kaminski, N.: Single-cell rna-seq reveals ectopic and aberrant lung-resident cell populations in idiopathic pulmonary fibrosis. Science Advances 6(28) (2020) 10.1126/sciadv.aba1983

[25] Valenzi, E., Bulik, M., Tabib, T., Morse, C., Sembrat, J., Trejo Bittar, H., Rojas, M., Lafyatis, R.: Single-cell analysis reveals fibroblast heterogeneity and myofibroblasts in systemic sclerosis-associated interstitial lung disease. Annals of the Rheumatic Diseases 78(10), 1379–1387 (2019) 10.1136/annrheumdis-2018-214865

[26] Pierce, E.M., Carpenter, K., Jakubzick, C., Kunkel, S.L., Evanoff, H., Flaherty, K.R., Martinez, F.J., Toews, G.B., Hogaboam, C.M.: Idiopathic pulmonary fibrosis fibroblasts migrate and proliferate to cc chemokine ligand 21. European Respiratory Journal 29(6), 1082–1093 (2007) 10.1183/09031936.00122806

[27] Pierce, E.M., Carpenter, K., Jakubzick, C., Kunkel, S.L., Flaherty, K.R., Martinez, F.J., Hogaboam, C.M.: Therapeutic targeting of cc ligand 21 or cc chemokine receptor 7 abrogates pulmonary fibrosis induced by the adoptive transfer of human pulmonary fibroblasts to immunodeficient mice. The American Journal of Pathology 170(4), 1152–1164 (2007) 10.2353/ajpath.2007.060649

[28] Mothes, R., Pascual-Reguant, A., Koehler, R., Liebeskind, J., Liebheit, A., Bauherr, S., Philipsen, L., Dittmayer, C., Laue, M., Manitius, R., Elezkurtaj, S., Durek, P., Heinrich, F., Heinz, G.A., Guerra, G.M., Obermayer, B., Meinhardt, J., Ihlow, J., Radke, J., Heppner, F.L., Enghard, P., Stockmann, H., Aschman, T., Schneider, J., Corman, V.M., Sander, L.E., Mashreghi, M.-F., Conrad, T., Hocke, A.C., Niesner, R.A., Radbruch, H., Hauser, A.E.: Distinct tissue niches direct lung immunopathology via ccl18 and ccl21 in severe covid-19. Nature Communications 14(1) (2023) 10.1038/s41467-023-36333-2

[29] Reyfman, P.A., Walter, J.M., Joshi, N., Anekalla, K.R., McQuattie-Pimentel, A.C., Chiu, S., Fernandez, R., Akbarpour, M., Chen, C.-I., Ren, Z., Verma, R., Abdala-Valencia, H., Nam, K., Chi, M., Han, S., Gonzalez-Gonzalez, F.J., Soberanes, S., Watanabe, S., Williams, K.J.N., Flozak, A.S., Nicholson, T.T., Morgan, V.K., Winter, D.R., Hinchcliff, M., Hrusch, C.L., Guzy, R.D., Bonham, C.A., Sperling, A.I., Bag, R., Hamanaka, R.B., Mutlu, G.M., Yeldandi, A.V., Marshall, S.A., Shilatifard, A., Amaral, L.A.N., Perlman, H., Sznajder, J.I., Argento, A.C., Gillespie, C.T., Dematte, J., Jain, M., Singer, B.D., Ridge, K.M., Lam, A.P., Bharat, A., Bhorade, S.M., Gottardi, C.J., Budinger, G.R.S., Misharin, A.V.: Single-cell transcriptomic analysis of human lung provides insights into the pathobiology of pulmonary fibrosis. American Journal of Respiratory and Critical Care Medicine 199(12), 1517–1536 (2019) 10.1164/rccm.201712-2410oc

[30] Morse, C., Tabib, T., Sembrat, J., Buschur, K.L., Bittar, H.T., Valenzi, E., Jiang, Y., Kass, D.J., Gibson, K., Chen, W., Mora, A., Benos, P.V., Rojas, M., Lafyatis, R.: Proliferating spp1/mertk-expressing macrophages in idiopathic pulmonary fibrosis. European Respiratory Journal 54(2), 1802441 (2019) 10.1183/13993003.02441-2018

[31] Mould, K.J., Moore, C.M., McManus, S.A., McCubbrey, A.L., McClendon, J.D., Griesmer, C.L., Henson, P.M., Janssen, W.J.: Airspace macrophages and monocytes exist in transcriptionally distinct subsets in healthy adults. American Journal of Respiratory and Critical Care Medicine 203(8), 946–956 (2021) 10.1164/rccm.202005-1989oc

[32] Büttner, M., Miao, Z., Wolf, F.A., Teichmann, S.A., Theis, F.J.: A test metric for assessing single-cell rna-seq batch correction. Nature Methods 16(1), 43–49 (2018) 10.1038/s41592-018-0254-1

[33] Zhang, X., Chen, Y., Hua, K., Xu, S., You, R., Hao, M., Li, W., Wei, L., Jia, J., Xi, X., Chen, S., Bian, H., Ye, M., Chen, A., Geng, Y., Liu, L., Luo, J., Fei, J., Lv, H., Zhang, P., Jiang, R.: uniheart: An ensemble atlas of cardiac cells provides multifaceted portraits of the human heart (2023) 10.21203/rs.3.rs-3215038/v1

[34] Netskar, H., Pfefferle, A., Goodridge, J.P., Sohlberg, E., Dufva, O., Teichmann, S.A., Brownlie, D., Michaëlsson, J., Marquardt, N., Clancy, T., Horowitz, A., Malmberg, K.-J.: Pan-cancer profiling of tumor-infiltrating natural killer cells through transcriptional reference mapping. Nature Immunology 25(8), 1445–1459 (2024) 10.1038/s41590-024-01884-z

[35] Herpelinck, T., Ory, L., Verbraeken, T., Nasello, G., Barzegari, M., Bolander, J., Luyten, F.P., Tylzanowski, P., Geris, L.: An integrated single-cell atlas of the limb skeleton from development through adulthood (2022) 10.1101/2022.03.14.484345

[36] He, Z., Dony, L., Fleck, J.S., Sza·lata, A., Li, K.X., Slišković, I., Lin, H.-C., Santel, M., Atamian, A., Quadrato, G., Sun, J., Pasca, S.P., Amin, N.D., Kelley, K.W., Bertucci, T., Temple, S., Bowles, K.R., Caporale, N., Villa, E., Testa, G., Cruceanu, C., Binder, E.B., Camp, J.G., Theis, F.J., Treutlein, B.: An integrated transcriptomic cell atlas of human neural organoids. Nature 635(8039), 690–698 (2024) 10.1038/s41586-024-08172-8

[37] Hrovatin, K., Bastidas-Ponce, A., Bakhti, M., Zappia, L., Büttner, M., Salinno, C., Sterr, M., Böttcher, A., Migliorini, A., Lickert, H., Theis, F.J.: Delineating mouse-cell identity during lifetime and in diabetes with a single cell atlas. Nature Metabolism 5(9), 1615–1637 (2023) 10.1038/s42255-023-00876-x

[38] Bandesh, K., Motakis, E., Nargund, S., Kursawe, R., Selvam, V., Bhuiyan, R.M., Eryilmaz, G.N., Krishnan, S.N., Spracklen, C.N., Ucar, D., Stitzel, M.L.: Single-cell decoding of human islet cell type-specific alterations in type 2 diabetes reveals converging genetic- and state-driven-cell gene expression defects (2025) 10.1101/2025.01.17.633590

[39] Jenkins, B.H., Tracy, I., Rodrigues, M.F.S.D., Smith, M.J.L., Martinez, B.R., Edmond, M., Mahadevan, S., Rao, A., Zong, H., Liu, K., Aggarwal, A., Li, L., Diehl, L., King, E.V., Bates, J.G., Hanley, C.J., Thomas, G.J.: Single cell and spatial analysis of immune-hot and immune-cold tumours identifies fibroblast subtypes associated with distinct immunological niches and positive immunotherapy response. Molecular Cancer 24(1) (2025) 10.1186/s12943-024-02191-9

[40] N. M., P., Fullard, J.F., Clarence, T., Mathur, D., Casey, C., Hennigan, E., Alvia, M., Krause-Massaguer, J., Barreda, A., Davis, D.A., Vontell, R.T., Garamszegi, S.P., Vance, J.M., Sang, L., Chatigny, M., Vismer, D., Landin, B., Burstein, D., Lee, D., Voloudakis, G., Berretta, S., Haroutunian, V., Scott, W.K., Bendl, J., Roussos, P.: A multi-region single nucleus transcriptomic atlas of parkinson’s disease. Scientific Data 11(1) (2024) 10.1038/s41597-024-04117-y

[41] Chen, M., Dahl, A.: A robust model for cell type-specific interindividual variation in single-cell rna sequencing data. Nature Communications 15(1) (2024) 10.1038/s41467-024-49242-9

[42] Yazar, S., Alquicira-Hernandez, J., Wing, K., Senabouth, A., Gordon, M.G., Andersen, S., Lu, Q., Rowson, A., Taylor, T.R.P., Clarke, L., Maccora, K., Chen, C., Cook, A.L., Ye, C.J., Fairfax, K.A., Hewitt, A.W., Powell, J.E.: Single-cell eqtl mapping identifies cell type–specific genetic control of autoimmune disease. Science 376(6589) (2022) 10.1126/science.abf3041

[43] Dann, E., Henderson, N.C., Teichmann, S.A., Morgan, M.D., Marioni, J.C.: Differential abundance testing on single-cell data using k-nearest neighbor graphs. Nature Biotechnology 40(2), 245–253 (2021) 10.1038/s41587-021-01033-z

[44] Peyré, G., Cuturi, M.: Computational Optimal Transport. now Publishers Inc, ??? (2019). 10.1561/9781680835519. http://dx.doi.org/10.1561/9781680835519

[45] Schiebinger, G., Shu, J., Tabaka, M., Cleary, B., Subramanian, V., Solomon, A., Gould, J., Liu, S., Lin, S., Berube, P., Lee, L., Chen, J., Brumbaugh, J., Rigollet, P., Hochedlinger, K., Jaenisch, R., Regev, A., Lander, E.S.: Optimal-transport analysis of single-cell gene expression identifies developmental trajectories in reprogramming. Cell 176(4), 928–94322 (2019) 10.1016/j.cell.2019.01.006

[46] Klein, D., Palla, G., Lange, M., Klein, M., Piran, Z., Gander, M., Meng-Papaxanthos, L., Sterr, M., Saber, L., Jing, C., Bastidas-Ponce, A., Cota, P., Tarquis-Medina, M., Parikh, S., Gold, I., Lickert, H., Bakhti, M., Nitzan, M., Cuturi, M., Theis, F.J.: Mapping cells through time and space with moscot. Nature 638(8052), 1065–1075 (2025) 10.1038/s41586-024-08453-2

[47] Székely, G.J., Rizzo, M.L.: Energy statistics: A class of statistics based on distances. Journal of Statistical Planning and Inference 143(8), 1249–1272 (2013) 10.1016/j.jspi.2013.03.018

[48] Peidli, S., Green, T.D., Shen, C., Gross, T., Min, J., Garda, S., Yuan, B., Schumacher, L.J., Taylor-King, J.P., Marks, D.S., Luna, A., Blüthgen, N., Sander, C.: scperturb: harmonized single-cell perturbation data. Nature Methods 21(3), 531–540 (2024) 10.1038/s41592-023-02144-y

[49] Novella-Rausell, C., Grudniewska, M., Peters, D.J.M., Mahfouz, A.: A comprehensive mouse kidney atlas enables rare cell population characterization and robust marker discovery. iScience 26(6), 106877 (2023) 10.1016/j.isci.2023.106877

[50] Swamy, V.S., Fufa, T.D., Hufnagel, R.B., McGaughey, D.M.: Building the mega single-cell transcriptome ocular meta-atlas. GigaScience 10(10) (2021) 10.1093/gigascience/giab061

[51] Wu, Y., Fan, Y., Miao, Y., Li, Y., Du, G., Chen, Z., Diao, J., Chen, Y.-A., Ye, M., You, R., Chen, A., Chen, Y., Li, W., Guo, W., Dong, J., Zhang, X., Wang, Y., Gu, J.: uniliver: a human liver cell atlas for data-driven cellular state mapping (2023) 10.1101/2023.12.09.570903

[52] Xu, C., Lopez, R., Mehlman, E., Regier, J., Jordan, M.I., Yosef, N.: Probabilistic harmonization and annotation of single-cell transcriptomics data with deep generative models. Molecular Systems Biology 17(1) (2021) 10.15252/msb.20209620

[53] Wolf, F.A., Angerer, P., Theis, F.J.: Scanpy: large-scale single-cell gene expression data analysis. Genome Biology 19(1) (2018) 10.1186/s13059-017-1382-0

[54] Traag, V.A., Waltman, L., Eck, N.J.: From louvain to leiden: guaranteeing well-connected communities. Scientific Reports 9(1) (2019) 10.1038/s41598-019-41695-z

[55] McInnes, L., Healy, J., Melville, J.: UMAP: Uniform Manifold Approximation and Projection for Dimension Reduction. arXiv (2018). 10.48550/ARXIV.1802.03426. https://arxiv.org/abs/1802.03426

[56] Dong, W., Moses, C., Li, K.: Efficient k-nearest neighbor graph construction for generic similarity measures. In: Proceedings of the 20th International Conference on World Wide Web. WWW ‘11. ACM, ??? (2011). 10.1145/1963405.1963487. http://dx.doi.org/10.1145/1963405.1963487

[57] Blondel, V.D., Guillaume, J.-L., Lambiotte, R., Lefebvre, E.: Fast unfolding of communities in large networks. Journal of Statistical Mechanics: Theory and Experiment 2008(10), 10008 (2008) 10.1088/1742-5468/2008/10/p10008

[58] Korsunsky, I., Millard, N., Fan, J., Slowikowski, K., Zhang, F., Wei, K., Baglaenko, Y., Brenner, M., Loh, P.-r., Raychaudhuri, S.: Fast, sensitive and accurate integration of single-cell data with harmony. Nature Methods 16(12), 1289–1296 (2019) 10.1038/s41592-019-0619-0

